# Elevated rates of positive selection drive the evolution of pestiferousness in the Colorado potato beetle (*Leptinotarsa decemlineata*, Say)

**DOI:** 10.1101/870543

**Authors:** Zachary P. Cohen, Kristian Brevik, Yolanda H. Chen, David J. Hawthorne, Benjamin D. Weibel, Sean D. Schoville

**Author notes:** Corresponding author: Zachary Cohen, Postal address: University of Wisconsin-Madison, Department of Entomology, 1630 Linden Drive, 637 Russell Labs, Madison, WI 53706, U.S.A.

## Abstract

Insect pests are characterized by expansion, preference and performance on agricultural crops, high fecundity and rapid adaptation to control methods, which we collectively refer to as pestiferousness. Which organismal traits and evolutionary processes facilitate certain taxa becoming pests remains an outstanding question for evolutionary biologists. In order to understand these features, we set out to test the relative importance of genomic properties that underlie the rapid evolution of pestiferousness in the emerging pest model: the Colorado potato beetle (CPB), *Leptinotarsa decemlineata* Say. Within the *Leptinotarsa* genus, only CPB has risen to pest status on cultivated *Solanum*. Using whole genomes from ten closely related *Leptinotarsa* species, we reconstructed a high-quality species tree of this genus. Within this phylogenetic framework, we tested the relative importance of four drivers of rapid adaptation: standing genetic variation, gene family expansion and contraction, transposable element variation, and protein evolution. Throughout approximately 20 million years of divergence, *Leptinotarsa* show little evidence of gene family turnover or transposable element variation contributing to pest evolution. However, there is a clear pattern of pest lineages experiencing greater rates of positive selection on protein coding genes, as well as retaining higher levels of standing genetic variation. We also identify a suite of positively selected genes unique to the Colorado potato beetle that are directly associated with pestiferousness. These genes are involved in xenobiotic detoxification, chemosensation, and hormones linked with pest behavior and physiology.

## Introduction

Among the many species that evolve rapidly, arthropods are among the most noteworthy in their ability to adapt to novel environments and pressures (Pélissié et al. 2018). Arthropod pests are responsible for billions of dollars of agricultural damage per year (Bebber 2015; Deutsch et al. 2018), and successful agricultural pests possess traits that allow them to 1) exploit agriculturally significant crops and 2) rapidly adapt to pesticides and other control tactics (Kim and McPheron 1993). Despite the considerable economic impact associated with these species, there is a lack of understanding as to how certain species evolve rapidly to agroecosystems (Chen and Schoville 2018). The organismal traits linked to pest success in agricultural environments remains open to debate, but could include pre-adaptive qualities such as: coevolution with chemically defensive plants, generalist herbivory behavior, high fecundity, high mutation rate, and short generation time (Dermauw et al. 2013; Cingel et al. 2016; Hardy et al. 2017; Dermauw et al. 2018). Additionally, due to contemporary genomic studies, certain features of species’ genomes appear advantageous to rapid evolution: elevated standing genetic diversity (Pearse et al., 2014; Lai et al., 2019), mutability linked to transposable elements (Baucom et al. 2009; Ayarpadikannan and Kim 2014; Pereira and Ryan 2019), gene family diversification (Hahn et al. 2007), and positive selection of regulatory and structural genes (Whitehead et al. 2017). Distinguishing the relative importance of these genomic properties, as well as whether they existed before “pest” status or after expansion to agroecosystems, requires a broad empirical foundation driven by comparative genomics research and phylogenetic understanding.

The Colorado potato beetle (CPB), *Leptinotarsa decemlineata* Say and related non-pest *Leptinotarsa* species, represent an ideal group to study the evolution of pestiferousness. CPB is one of the best studied insect species on the planet with more than 10,000 research articles documenting its biology, including a comprehensive annotated reference genome (Schoville et al. 2018). *Leptinotarsa* Chevrolat (Coleoptera: Chrysomelidae) is a phytophagous genus of beetle that includes approximately 41 species (Jacques 1988). Although the *Leptinotarsa* beetles have similar ecological niches, overlapping geographical ranges, and host plant specificity, only CPB is recognized as a major agricultural pest. CPB is able to utilize at least a dozen plants in the family Solanaceae as suitable hosts, it has the greatest host breadth, and has repeatedly expanded onto several agriculturally significant crops including potato, tomato, eggplant, pepper and tobacco (Hsiao and Fraenkel, 1968b; Duchesne and Parent, 1991; Jacques, 1988). While several other members of the *Leptinotarsa* genus also use multiple host plants, such as *L. haldemani*, (Rogers), *L. rubiginosa*, (Jacoby), and *L. peninsularis* (Horn) (Jacques 1988), none have become a significant pest. In addition to exploiting agricultural crops, CPB has demonstrated an extraordinary ability to rapidly evolve resistance to diverse insecticides, now exceeding more than 50 chemical products across all major modes of action (Casagrande 1987; Alyokhin et al. 2008). Indeed, the beetle ranks among the fastest insect pests to evolve resistance to new insecticides (Brevik et al. 2018). Additionally, this beetle has readily invaded and established in challenging environments where potato is now grown, such as Northern Europe (Grapputo et al. 2005), adjusting its life history to cope with cold winters and short photoperiods. In contrast, other *Leptinotarsa* species are restricted to more temperate regions (Hiiesaar et al. 2016).

Upon publishing the genome of CPB, Schoville *et al*. (2018) raised two hypotheses for genomic properties that could enable rapid evolution. The first, based on observed patterns of single nucleotide polymorphism in the CPB genome, was that the high standing genetic diversity could facilitate rapid evolutionary change. The second proposed mechanism was that transposable elements, which comprised >17% of the genome and appear to be rapidly evolving relative to other Coleoptera genomes, could be a source of increased genetic novelty. Here, we apply a comparative genomics framework using nine additional *Leptinotarsa* species to test these hypotheses as well as two additional mechanisms of rapid evolution: gene family diversification and selection on regulatory and structural genes (see our hypothesis testing framework in **Table 1**). We directly test for an association between these genomic drivers of adaptation in the *Leptinotarsa* genomes by incorporating their evolutionary history, via a robust phylogeny, and determine if these features differ between pest and non-pest individuals within the CPB lineage.

**Table 1.** Hypothesis testing framework for genomic features that enable the rapid evolution of pestiferousness.

| Genomic feature | Species level: CPB with related *Leptinotarsa* species (pre-adaptive)<br>Prediction | CPB level: shared among non-pest and pest CPB populations (pre-adaptive)<br>Prediction | CPB level: pest CPB populations only (adaptive)<br>Prediction |
| --- | --- | --- | --- |
| Transposable elements | Significant and strong phylogenetic signal. | Significant elevation of percent of CPB genome. | Significant elevation of percent of pest CPB genome. |
| Gene family diversification | Significant changes in gene family size among all species at phenotypically relevant families | Significant changes in gene family size associated with host plant detection, chemosensation, and xenobiotic metabolism at CPB | Significant changes in gene family size associated with host plant detection, chemosensation, and xenobiotic metabolism at only pest CPB |
| Positive selection on regulatory and structural genes | Many significant genes associated with species success shared among closely related species and/or total rate of significant genes explained phylogenetically. | Many significant genes associated with species success and/or elevated rates of selection at the phylogenetic level. | Many significant genes associated with species success and/or elevated rates of selection at the phylogenetic level |
| Genetic diversity | Significant and strong phylogenetic signal. | Greatest in CPB lineage, indicative of large effective population size | Reduced in CPB lineage, indicative of founder events and intensive selection in agricultural context. |

## Materials and Methods

### Sampling and Genome Sequencing

We collected nine non-pest *Leptinotarsa* species from natural populations in North America: *L. juncta* Germar, *L. haldemani* Rogers, *L. lineolata* Stål, *L. peninsularis* Horn, *L. rubiginosa* Rogers, *L. texana* Schaeffer, *L. tumamoca* Tower, and *L. undecemlineata* (see **Table 2** for detailed collection information). For *L. decemlineata* (CPB), we used three individuals: the published reference genome, which represents a beetle from a population in Long Island, New York with a long history of developing resistance to insecticides (Schoville et al., 2018), as well as two additional individuals. One non-pest individual was collected in Kansas on its natal host plant, Buffalo bur, *Solanum rostratum* Dunal, with no prior exposure to insecticides, and an additional pest specimen was collected in Wisconsin from a beetle population in a conventional potato field known to be resistant to neonicotinoids. Furthermore, beetle populations on Buffalo bur are considered ancestral to the potato-feeding pest populations (Izzo et al. 2018). For each individual, genomic DNA was isolated using DNEasy Blood & Tissue Kits (Qiagen, Hilden, Germany), according to the manufacturer’s protocols. The UW Biotech Center constructed sequencing libraries optimized for 150bp paired end reads and each sample was sequenced to approximately 20x coverage on an Illumina HiSeq2500. These data are available at NCBI on Genbank (Bioproject PRJNA580490, short read archive accessions SRR10388401:SRR10388312).

**Table 2.** Genome assembly statistics for *Leptinotarsa* species.

| Species | Collecting location | N50 (bp) | Total size (bp) | BUSCO from 2442 Single Copy Orthologs | Coverage |
| --- | --- | --- | --- | --- | --- |
| *L. decemlineata* | Long Island NY USA | 139401 | 642052436 | 97.4% [C:91.6%, F:5.8%] | 140.0 LI<sup>1</sup> |
| *L. decemlineata* | Hays KS USA | 139401 | 642052436 | 97.4% [C:91.6%, F:5.8%] | 6.49 |
| *L. decemlineata* | Middleton WI USA | 139401 | 642052436 | 97.4% [C:91.6%, F:5.8%] | 5.10 |
| *L. lineolata* | Pima Co Arizona USA | 6592 | 643141069 | 85.8% [C:54.7%, F:31.1%] | 19.0 |
| *L. texana* | Hidalgo Texas USA | 6003 | 636866419 | 85.4% [C:51.6%, F:33.8%] | 18.9 |
| *L. undecemlineata* | Chiapas Mexico | 5968 | 635369829 | 87.5% [C:56.4%, F:31.1%] | 21.1 |
| *L. defecta* | Rio Grande City Texas USA | 5572 | 618519877 | 87.6% [C:55.5%, F:32.1%] | 19.2 |
| *L. tumamoca* | Pima Co. Arizona USA | 4261 | 570462719 | 85.4% [C:52.7%, F:32.7%] | 17.7 |
| *L. peninsularis* | Pima Co. Arizona USA | 3890 | 549858680 | 85.3% [C:50.8%, F:34.5%] | 16.8 |
| *L. juncta* | Gainesville FL USA | 2990 | 512812428 | 79.8% [C:41.2%, F:38.6%] | 20.3 |
| *L. rubiginosa* | Yavapai Co. Arizona USA | 2928 | 558153188 | 80.8% [C:43.5%, F:37.3%] | 19.3 |
| *L. haldemani* | Yavapai Co. Arizona USA | 2571 | 533712537 | 71% [C:38.4%, F:32.6%] | 22.6 |

### Whole Genome Assembly and Assessment

For each of the nine additional *Leptinotarsa* species, the raw reads were pre-processed for quality and read length using the programming suite BBmap v36.30 (Bushnell 2014). These curated reads were used for whole genome assembly in SPAdes (St. Petersburg genome assembler) v3.9.0 (Bankevich et al. 2012). Genome assembly of the nine non-pest species was run on the University of Wisconsin’s Center for High Throughput computing (CHTC http://chtc.cs.wisc.edu/). Furthermore, as the data were obtained from field collected individuals, they were assumed to be highly heterozygous and *de novo* contigs were processed using Redundans v0.12c, with standard parameters for moderate coverage and highly heterozygous samples (Pryszcz and Gabald 2016). Genome assemblies are available at NCBI on Genbank (Bioproject PRJNA580490, whole genome shotgun accessions WOGI00000000:WOGR00000000).

Metrics of the cleaned assemblies were calculated using QUAST (Quality Assessment Tool for Genome Assemblies, http://quast.sourceforge.net/quast) and custom bash scripts. We assessed the completeness and quality of the draft assemblies using BUSCO v3 (Benchmarking Universal Single-Copy Orthologs, https://busco.ezlab.org/), with the Arthropoda Endopterygota v9 single copy ortholog set of 2,358 genes (Simao et al. 2015). Additionally, we used the genome assembly of the Asian longhorn beetle, *Anoplophora glabripennis* Motschulsky (Coleoptera: Cerambycidae), as an outgroup comparison for BUSCO results.

### Phylogenetic Reconstruction

The single copy orthologs identified in BUSCO were curated from each genome assembly to develop a data matrix for phylogenetic analysis. Of the 2,385 single-copy nuclear genes in the Endopterygota v9 dataset, approximately 1,510 sequences were shared among the ten *Leptinotarsa* species and the *A. glabripennis* outgroup. Alignments of these single-copy genes were generated using the spALN program (Gotoh 2008), which searches genome assemblies for shared orthologs. Isolated orthologs for each species were then aligned to each other using a custom C program CATaNNN v1.0 (Weibel and Cohen 2018) and MAFFT (Katoh et al. 2002). GUIDANCE2 (Penn et al. 2010) was used to curate the alignments for quality. This resulted in a final set of 1,418 single copy orthologue alignments that were used for phylogenetic reconstruction. A gene tree was estimated for each of the 1,418 gene alignment in RAxML v8.2.10 (Stamatakis 2014), assuming a generalized time-reversible, GTR + γ substitution model, and support values were generated using 100 bootstraps with 20 maximum likelihood searches under a gamma distribution (Tavaré 1986; Stamatakis 2014). The resulting 1,418 trees were then used to jointly estimate a species tree under a multi-species coalescent model via ASTRAL-II v5 (Mirarab et al. 2014). This method infers a species phylogeny given the input gene trees, where branch lengths are measured at internal nodes in terms of coalescent units, and support for each quartet is calculated as a local posterior probability value. To estimate the divergence of different species in relative time, we first rescaled the branch lengths using substitutions per site in RaXML. Second, we time calibrated the tree using the program r8s v1.81 (Sanderson 2003), based on the previous estimate of the divergence time between *L. decemlineata* and *A. glabripennis* of 104.5 Ma (Thomas et al. 2018).

### Estimating Levels of Standing Genetic Diversity

To estimate standing genetic diversity, we calculated a genome-wide measure of heterozygosity, *h*, for each species using ANGSD (Korneliussen et al. 2014). Individual heterozygosity (conversely, homozygosity) is a measure of genetic polymorphism that is dependent on the effective population size and inbreeding patterns in a population (Gregorius 1978). Levels of polymorphism can also be reduced by episodes of population size contraction and/or selection (Nei et al. 1975; Tajima 1989). CPB has a history of population outbreaks, with some exceptional numbers observed after it expanded onto cultivated potato. Such events were followed by chemical control efforts that the beetle rapidly adapted to. The consequences of these boom and bust periods on the pest population heterozygosity may or may not be evident (Crossley et al. 2017), but standing genetic variation in non-pest individuals should be similar to close relatives. Thus, we examined different CPB individuals (pest and non-pest) to ensure levels of polymorphism in this species are not biased by the invasion history (Izzo et al. 2018) or numerous episodes of selection from pesticide exposure.

### Expansion of Transposable Elements

In order to determine if rapid adaptation could be associated with transposable element (TE) variation, we utilized two specialized programs to identify, curate, and quantify TE abundance among the ten *Leptinotarsa* species. We used RepeatMasker v4.0 (Smit et al., 2015) to find known TEs from Repbase (Arkhipova et al. 2012), and RepeatModeler (Smit and Hubley 2008), which detects *de novo* repeat elements. We also performed extensive literature searches to locate novel transposons that were not within Repbase. All three libraries were subsequently used within RepeatMasker to determine the overall TE content and percent of total genome in the ten *Leptinotarsa* species. The quantities for TE groups were then tested for significant changes in CAFE v4.2 (De Bie et al. 2006).

### Gene Family Evolution

Gene family evolution involves either a significant contraction or expansion in the number of protein coding genes, which may change the molecular diversity or biological function of the family (Olson 1999; Schrider and Hahn 2010). Expansion and contraction of gene families among extant and ancestral lineages of the *Leptinotarsa* (quantified as the birth-death parameter, lambda λ) was estimated while accounting for phylogenetic history, using the software CAFE v4.2 (De Bie et al. 2006). Gene families were classified as orthogroups based on inferred homology using the complete Arthropoda Orthodb v8 dataset of 38,195 known orthogroups (Waterhouse et al. 2013). Paralogs from this set were identified in the draft genomes of the ten *Leptinotarsa* species using the spALN program (Gotoh 2008). Given the fragmentation of these shotgun genomes, we controlled the redundancy of identified orthologs by removing 1) matches to non-insect taxa in the data set, 2) de-duplicating redundant sequences, based on percent similarity (<10%) and a minimum overlap of four sequences of at least 31 bases, using the dedupe program of the BBMAP suite, and 3) isolating loci based on unique scaffold location, *i.e*. no paralogs were included that shared the same position in the genomes. The time calibrated species tree and number of genes per orthogroup were used to estimate λ. Additionally, error modelling and a significance threshold of 0.01 were used in CAFE to reduce type I error. Significantly variable orthogroups were then associated with known gene ontology (GO) terms and cross referenced with significantly enriched GO terms from the branch-site results.

### Positive Selection in Protein Coding Genes

The evolution of protein coding genes among closely related species was examined for signatures of positive selection in the form of elevated non-synonymous substitutions (Yang and Nielsen 2002). We conducted a genome-wide analysis of protein coding genes using the community annotated official gene set (OGS v1.1) of the Colorado potato beetle (Schoville et al. 2018). This gene set consists of 24,853 genes and was used to identify shared orthologs in each *Leptinotarsa* genome using the spALN program (Gotoh 2008). Putative coding regions in each species were curated for their likelihood of being a true ortholog, based on the best reciprocal hit with respect to the reference CPB sequence, their position in each species’ assembly, and their percent coverage. Subsequently, curated orthologs from the nine other species were then aligned to the reference CPB genes, to generate alignments as described previously for the single copy ortholog data set, and again curated for the three main parameters associated with quality and degree of certainty, as outlined by Schneider *et al*. (2009).

Due to the fragmentation of the assemblies, as well as possible incorporation of paralogs, we corrected for mis-alignment and quality using Guidance v2.02, at high stringency, with 30 bootstraps per alignment (Penn et al. 2010). To avoid spurious codon substitutions introduced by assembly errors, we replaced ambiguous alignments (including internal stop codons and frameshift mutations) with N’s using the program MACSE (Ranwez et al. 2018), and terminal stop codons were removed using a custom bash program. These curation steps resulted in a final set of gene alignments, which were used to test for positive selection among *Leptinotarsa* species via the adaptive branch-site random effects model (Smith et al. 2015).

The branch-site test determines locus specific positive selection among species within a phylogeny by detecting elevated rates of non-synonymous to synonymous (d_N_/d_s_) substitutions in protein-coding genes, while accounting for rate variation among the lineages (Smith et al. 2015). Thus, the method provides evidence for positive selection at each locus and on each branch of the species tree, rooted at *L. lineolata*. In order to determine the effects of individual sampling within the CPB branch, the branch-site test was run three separate times with CPB data alternatively represented by a pest (New York or Wisconsin) or non-pest (Kansas) individual. Putatively selected genes were subsequently grouped by GO terms using the *Leptinotarsa decemlineata* official gene set and the Gene Ontology project v1.2 database (Schoville et al., 2018; Ashburner et al. 2011; 2019). Loci under positive selection were curated by taxon and compared for similarity. These significant loci were further examined by gene ontology term and tested for enrichment.

Enrichment of GO terms of all significant loci among all species was determined via a one-way Fischer’s exact test. Shared and unique enriched terms of significant genes were assigned to lineages to make comparisons to the CPB branch. In order to resolve how genes associated with xenobiotic metabolism and insecticide resistance (detoxification enzymes, membrane transporters, putative binding sites, cuticular proteins, and sequestration proteins) might be changing among these beetles, a subset of 673 genes identified in Crossley *et al*. (2017) was further scrutinized for selection. The shared loci among the CPB individuals were examined again for codon-based positive selection using the Bayes empirical Bayes model of evolution from the PAML package (Yang et al. 2005).

### Testing for an Association between Genomic Features and Pestiferousness

In order to directly compare the four hypotheses of genomic properties that might drive rapid evolution and pestiferousness in the *Leptinotarsa* genus, we employed statistical tests that account for evolutionary history by referencing species relatedness and genetic distance (Felsenstein, 1985; Revell et al., 2008). Phylogenetic corrections are already employed in CAFE and the adaptive branch-site test. However, we were specifically interested in whether outstanding genomic features varied among taxa in a phylogenetic-dependent fashion (Jolivet et al. 1988; Garland 1992). In addition to genomic features, we investigated whether ecological host breadth might be explained by phylogenetic history and/or genomic features associated with adaptation, by incorporating an existing scale of host breadth based on Hsiao (Jolivet et al. 1988). Species were assigned categorical scores as follows: 1 = a specialist on one plant genus, 2 = oligophagous on more than one plant genus, but less than three genera, and 3 = a generalist on more than three genera. Phylogenetic independent contrast (PIC) values were determined for these features to account for trait evolution (Felsenstein 1985; Garland 1992). Here, we investigate 1) if any of these putatively adaptive features have a phylogenetic signal (*i.e*. whether or not a trait value is explained by evolutionary history) (Revell et al. 2008), 2) if particular lineages have the greatest phylogenetic signal, 3) if traits are evolving at different rates across the lineage, and 4) after accounting for non-independence, if the genomic features are correlated with one another. Using the R package phyloSignal v1.2.1 (Keck et al., 2016), we corrected the trait values for evolutionary dependency by generating PIC values for: total proportion of loci under selection, loci under selection associated with insecticide resistance, host plant breadth and heterozygosity for each lineage. These values were tested for phylogenetic dependency using Moran’s I and Blomberg’s K, which provide a local indicator of phylogenetic association (LIPA) and evolution, respectively (Keck et al. 2016).

In order to compare rates of trait evolution, we calculated standardized independent contrast values, which are PIC values corrected by the square root of branch length sums or the standard deviation, according to Garland (1992). We used an outlier test to determine the standardized rate values that differed the most, which allowed us to identify if trait evolution accelerated along a particular lineage (Garland 1992). Lastly, we compared independent contrast values for correlations to each other for total and candidate resistant loci proportions, host breadth, and heterozygosity in the R package analysis of phylogenetic evolution (APE) v5.3 (Paradis 2012).

## Results

### Assessment of Genome Assemblies, Species Tree and Heterozygosity

Genome assemblies of the ten *Leptinotarsa* species have highly similar total genome size (ranging from 520-650 MB; **Table 2**), with all assemblies having approximately 20-fold depth of coverage. BUSCO scores assessing completeness of the genomes vary from 71% (*L. haldemani*) to 97% (*L. decemlineata*), with the remaining species having BUSCO scores >80%. The time calibrated *Leptinotarsa* phylogeny of the ten species, with *A. glabripennis* as an outgroup, suggests divergence of this genus at around 19 million years ago (Ma) with the clade including CPB arising ~10 Ma (**Figure 1, Supplemental Figure S1**). Heterozygosity among *Leptinotarsa* species show that more recently diverged clades, which includes *L. decemlineata* (non-pest: KS: h=1.52%; pest: WI: h=1.45%; LI: h=0.91%)*, L. peninsularis* (h=1.77%) and *L. tumamoca* (h=1.75%), have high genome-wide heterozygosity and a signal of autocorrelation (p<0.05) (**Figure 2, Supplemental Figure S2**). However, this signal is lost when using the heterozygosity values for the pest CPB individuals, suggesting that the large heterozygosity for these species is associated with their position in the phylogeny and may contribute to the ability of CPB to become a successful pest.

**Figure 1.**
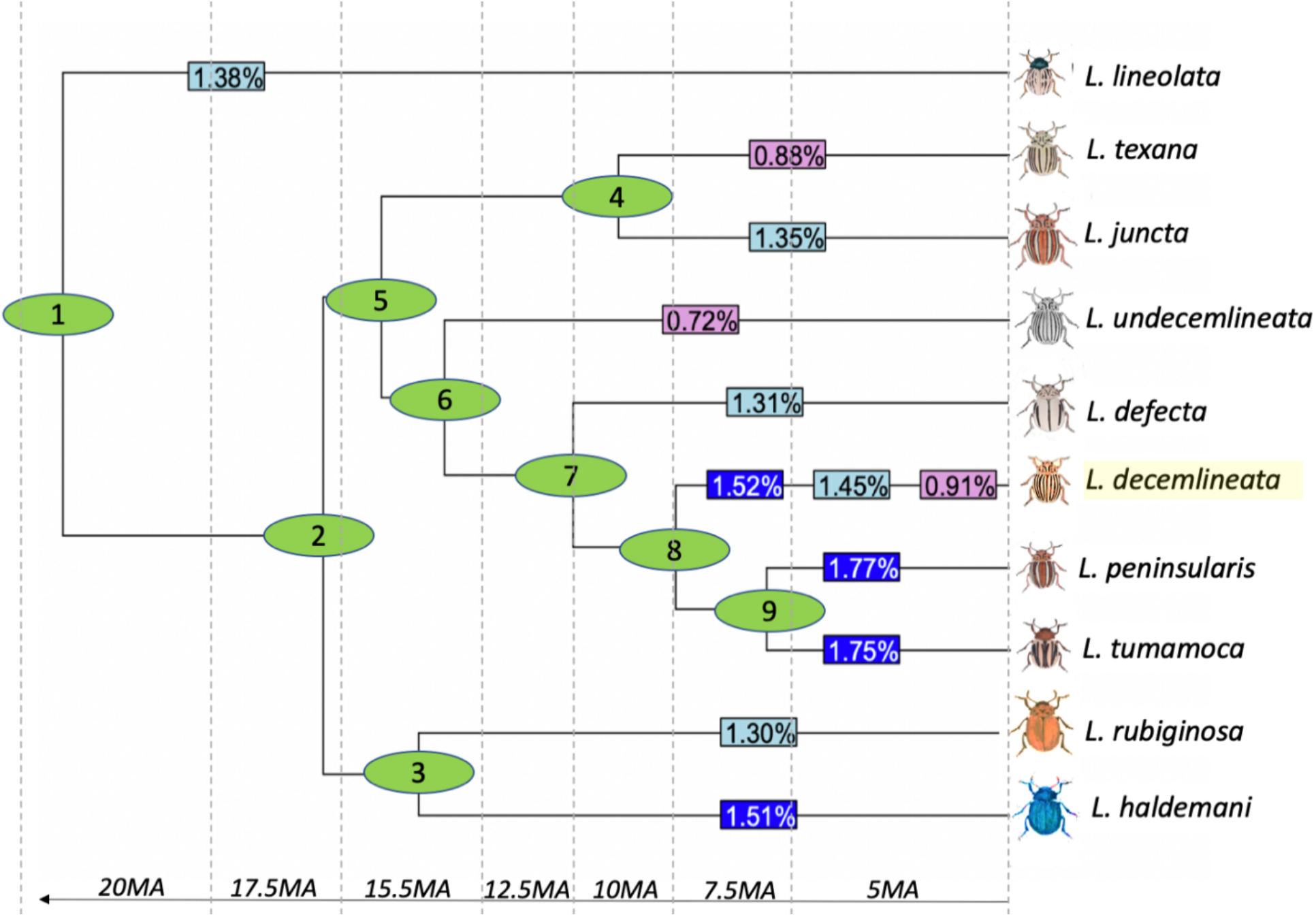
Ultrametric species tree of *Leptinotarsa* clade, rooted with *Anoplophora glabripennis* outgroup (not shown). Divergence time is stated as millions of years ago (Ma) and nodes are numbered for reference. Heterozygosity estimates are shown in boxes on branches, with color reflecting low (violet), moderate (light blue) or high (dark blue) levels of heterozygosity. Three conspecific CPB replicates are shown on the *L. decemlineata* branch as colored boxes. Left to right: non-pest Kansas individual, Wisconsin pest-resistant individual and Long Island pest-resistant individual.

**Figure 2.**
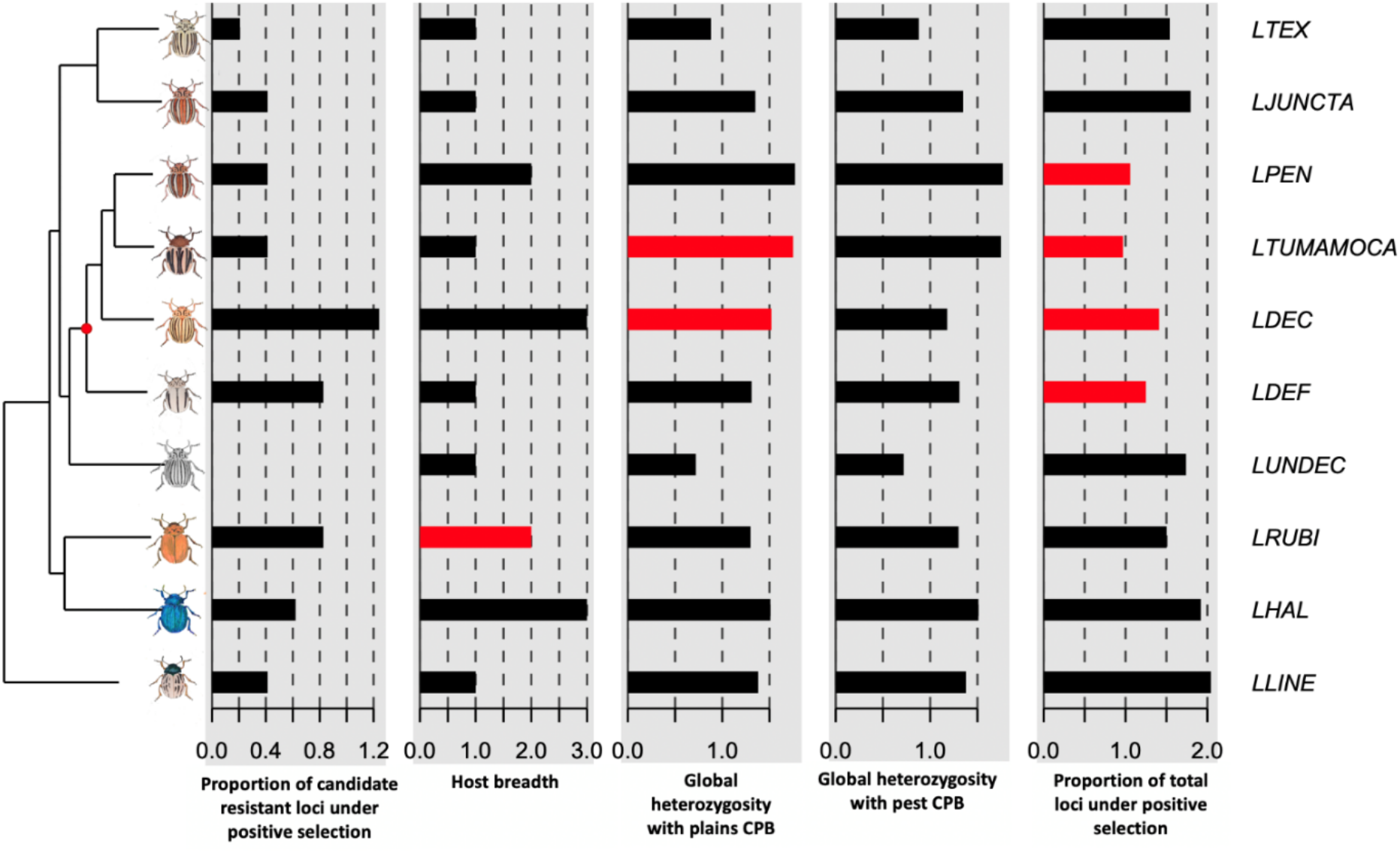
Local indicator of phylogenetic association (LIPA) based on Moran’s I for *Leptinotarsa* genomic features. Significant bars (p < 0.05) are shown in red.

### Gene Family Evolution

With respect to gene family size changes, and corresponding gene ontology annotations among *Leptinotarsa*, we find significantly more contractions than expansions (**Supplemental Figure S3, Supplemental Figure S4**). Of the 38,195 orthogroups (EOGS), 18,412 were represented in the *Leptinotarsa* genus. 37 EOGs with significant λ values (p ≤ 0.01) were identified among these species. The majority of unique EOGs expanding or contracting did not have GO terms that were enriched from the Fisher exact test. Within the twelve expanding EOGs, there were seventeen GO terms. Ten of these were shared with other expanding or contracting EOGs, and two of these were enriched for positive selection (GO:0003676: nucleic acid binding; GO:0003677: DNA binding). The seven remaining GO terms were uniquely expanding with an additional three GO terms significantly enriched (GO:0000166: nucleotide binding; GO:0005524: ATP binding; GO:0016887: ATPase activity) (**Supplemental Table S1**). Of the twenty-seven uniquely contracting orthogroups, five GO terms were enriched (GO:0003700: sequence-specific DNA binding transcription factor activity; GO:0006396: RNA processing; GO:0008270: zinc ion binding; GO:0016021: integral component of membrane; GO:0043565: sequence-specific DNA binding) (**Supplemental Table S1**).

At the CPB branch, only one term was contracting and is associated with cysteine-type peptidase activity. This gene ontology was not enriched in any lineage. *L. peninsularis* and *L. tumamoca* have the most contracting EOGs, 13 and seven respectively.

### Expansion of Transposable Elements

We found similar percentages of genome level transposable elements among the ten *Leptinotarsa* species investigated (**Supplemental Table S2**). Furthermore, there were only two transposable element groups experiencing significant copy number size changes among the species (**Supplemental Figure S5**, **Supplemental Tables S3**). The Gypsy-Cigr TE family was expanding in *L. decemlineata*, while also contracting in *L. juncta*. The tRNA TE family was contracting in *L. decemlineata*, while also expanding in *L. haldemani* and *L. tumamoca*.

### Positive Selection in Protein Coding Genes

The representation of the 24,826 genes from the CPB official gene set varied slightly among the three alignment sets with Long Island (18,749 of 24,826, ~75%) and Kansas (18,166 of 24,826, ~73%) having greater percentage of the OGS than the Wisconsin CPB alignments (17,237 of 24,826, ~69%). Loci under positive selection by species were compared for commonality (**Supplemental Figure S6**). The GO terms for the significant loci for each taxon and at each internal node in the *Leptinotarsa* phylogeny (**Table 3**), were tested for enrichment via a Fisher exact test. Among non-pest lineages, there was some overlap with respect to enriched GO terms and orthogroups significantly varying in size across the phylogeny. Fibrillarin component of small nuclear ribonucleoproteins in ribosomal RNA processing (GO:0003723) is enriched in all of the lineages except for *L. rubiginosa* yet was detected as an expanding gene family in *L. rubiginosa* and *L. texana* (**Supplemental Table S1**). The enriched GO term 0006508 (proteolysis) in *L. texana* is associated with the previously significant gene family expansion in EOG8M945N. Genes with a zinc finger domain (GO:0003676) were enriched in all taxa and are a significantly contracting orthogroup (EOG8P2SH4) in the *L. decemlineata* lineage.

**Table 3.**
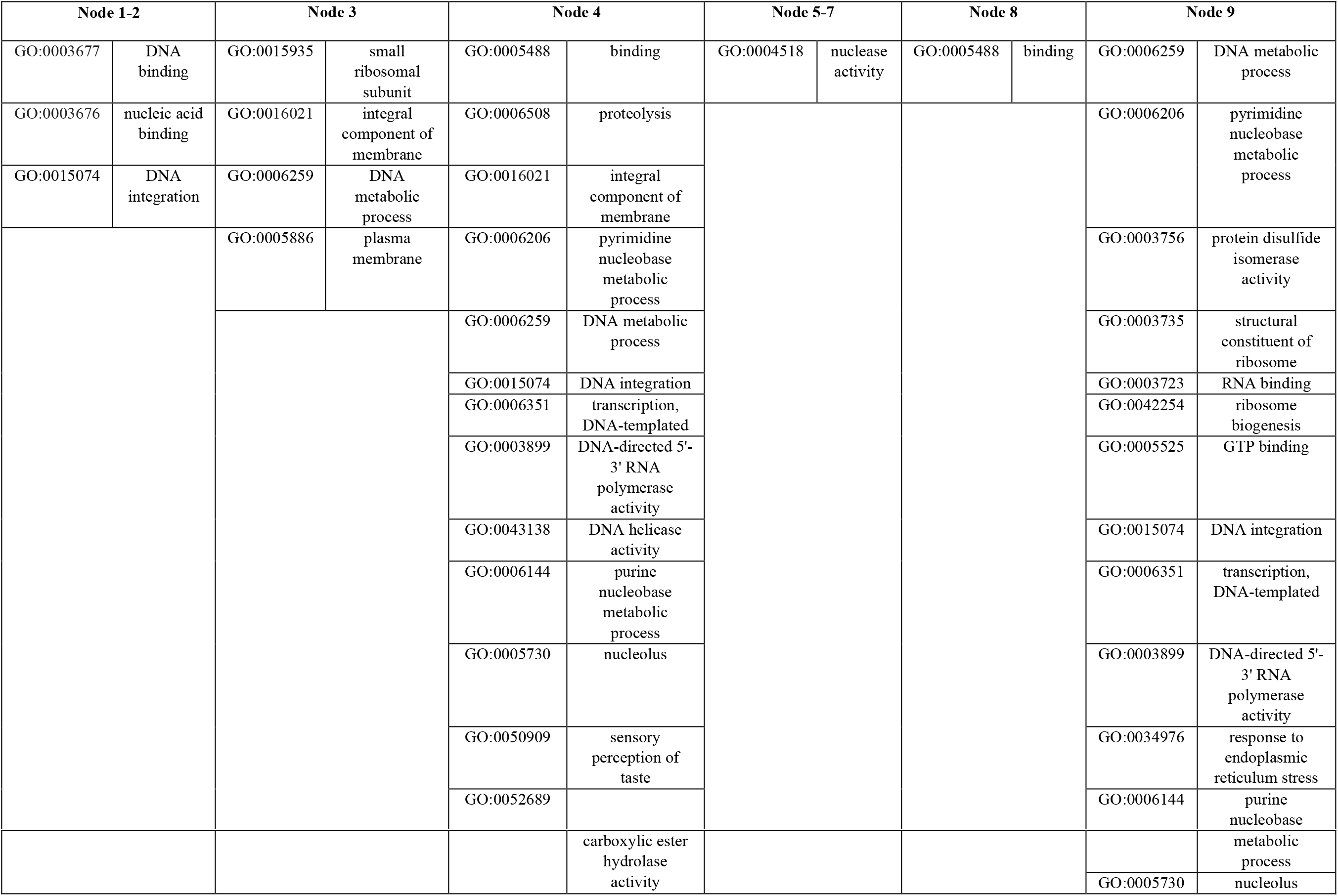
Shared GO term annotations represented by candidate positive selection genes at different nodes in the *Leptinotarsa* phylogeny.

Within the CPB lineage, there are shared and unique significant genes and enriched GO terms (**Supplemental Table S4, Supplemental Table S5, Supplemental Figure S7, Table 4**). The shared GO terms of the pest and non-pest CPB individuals are associated with sugar metabolism, neuronal development and signaling, and larval development (**Table 4**). Additionally, there were nine GO terms unique to the pest CPB (**Table 4**). More than half of the significant genes unique to the pest individuals were associated with enriched term: integral membrane component (GO:0016021). This GO includes genes annotated with xenobiotic metabolism (esterases, cytochrome P-450s, ATP-transporters etc.), chemosensation (gustatory receptors), hormonal receptors, and membrane-bound ligands. Other pest-specific terms included superoxide dismutase activity (GO:0004784) and removal of superoxide radicals (GO:0019430), the G protein-coupled receptor signaling pathway (GO:0007186 and 0004930), receptor (GO:0005057) and synapse signaling (GO:0045202), L-proline biosynthetic process (GO:0055129) and tracheal regulation (GO:0035159).

**Table 4.** Unique GO terms of selected genes within the *Leptinotarsa decemlineata* lineage: shared pest and non-pest (KS, LI, WI) and pest only (WI, LI).

| Combined pest and non-pest *Leptinotarsa decemlineata* GO terms and annotations |  | Shared Pest *Leptinotarsa decemlineata* GO terms and annotations |  |
| --- | --- | --- | --- |
| GO:0051963 | regulation of synapse assembly | GO:0004784 | superoxide dismutase activity |
| GO:0008376 | acetylgalactosaminyltransferase activity | GO:0016021 | integral component of membrane |
| GO:0004793 | threonine aldolase activity | GO:0035159 | regulation of tube length, open tracheal system |
| GO:0016339 | calcium-dependent cell-cell adhesion via plasma membrane cell adhesion molecules | GO:0007186 | G protein-coupled receptor signaling pathway |
| GO:0030111 | regulation of Wnt signaling pathway | GO:0004930 | G protein-coupled receptor activity |
| GO:0007367 | segment polarity determination | GO:0005057* | obsolete signal transducer activity, downstream of receptor |
| GO:0045773 | positive regulation of axon extension | GO:0055129 | L-proline biosynthetic process |
| GO:0045746 | negative regulation of Notch signaling pathway | GO:0019430 | removal of superoxide radicals |
| GO:0070593 | dendrite self-avoidance | GO:0045202 | synapse |
| GO:0048057 | R3/R4 development |  |  |
| GO:0005911 | cell-cell junction |  |  |

When putative insecticide resistant genes are grouped together by function, the percentage of genes with these functions was significantly greater in the two-pest resistant Colorado potato beetles than in other *Leptinotarsa* and the non-pest susceptible CPB (p-value = 0.0196) The four positively selected insecticide resistant loci shared among the three *L. decemlineata* individuals are a cytochrome P450, a ryanodine calcium channel, a glutamate chloride channel, and an esterase (**Table 5**). Codon selection analysis via the BEB test identified shared sites among all three CPB individuals, but notably additional significantly selected codons unique to the pest CPB samples (**Figure 3, Figure 4**).

**Table 5.** Genes associated with xenobiotic detoxification pathways in *Leptinotarsa decemlineata*.

| CPB-KS (S) |  | CPB- LI (R) |  | CPB-WI (R) |  |
| --- | --- | --- | --- | --- | --- |
| LDEC004414 | cuticular protein 4 | LDEC002144 | atp-binding cassette sub-family b member mitochondrial-like | LDEC011039 | cuticle protein cuticle |
| LDEC006847 | carboxylesterase 1E | LDEC005020 | cuticular protein analogous to peritrophins 1-b | LDEC023090 | esterase |
| LDEC017668 | atp-binding cassette sub-family g member 5 | LDEC005242 | larval cuticle protein 1 |  |  |
| CPB- LI + CPB-WI (R) |  | CPB-WI + LI+ KS (S+R) |  |  |  |
| LDEC011604 | cytochrome p450 6k1-like CYP | LDEC005267 | venom carboxylesterase-6 esterase |  |  |
| LDEC014122 | Esterase e4-like | LDEC007559 | glutamate receptor kainate 2-like glutamate chloride channel |  |  |
|  |  | LDEC016278 | probable cytochrome p450 305a1 CYP |  |  |
|  |  | LDEC018687 | voltage-dependent calcium channel gamma-7 subunit [Cephus cinctus] ryanodine.calcium.channel" |  |  |

**Figure 3.**
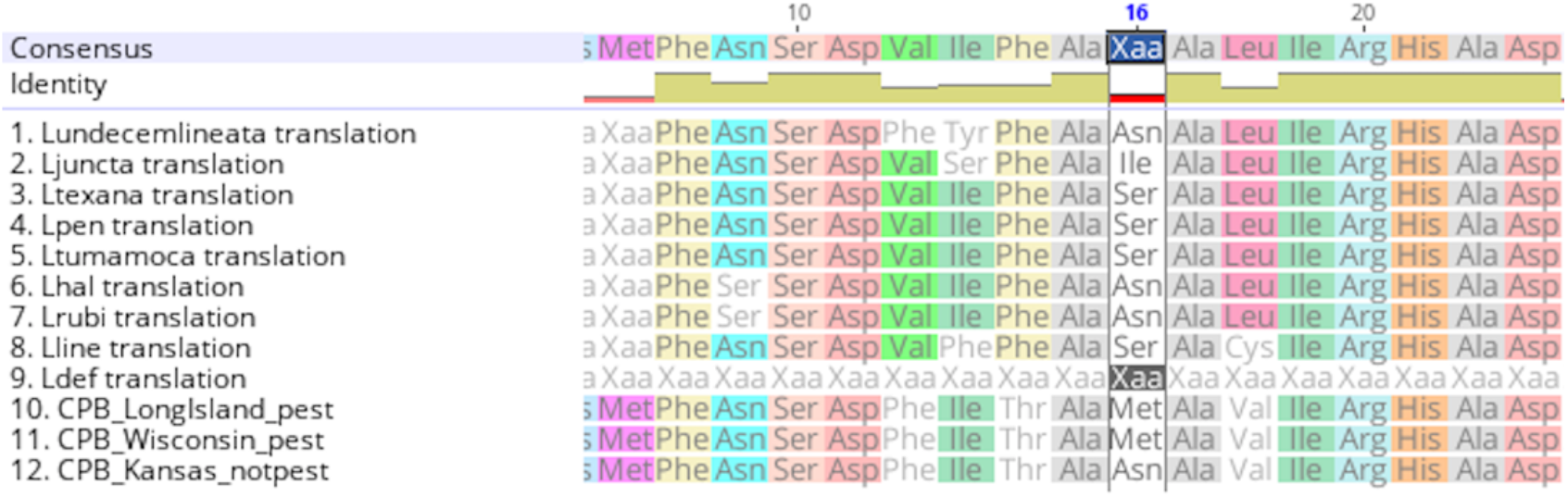
Candidate selection gene LDEC0014122, an esterase, shows significant codon change at position 16 in resistant CPB individuals (rows 10 and 11).

**Figure 4.**
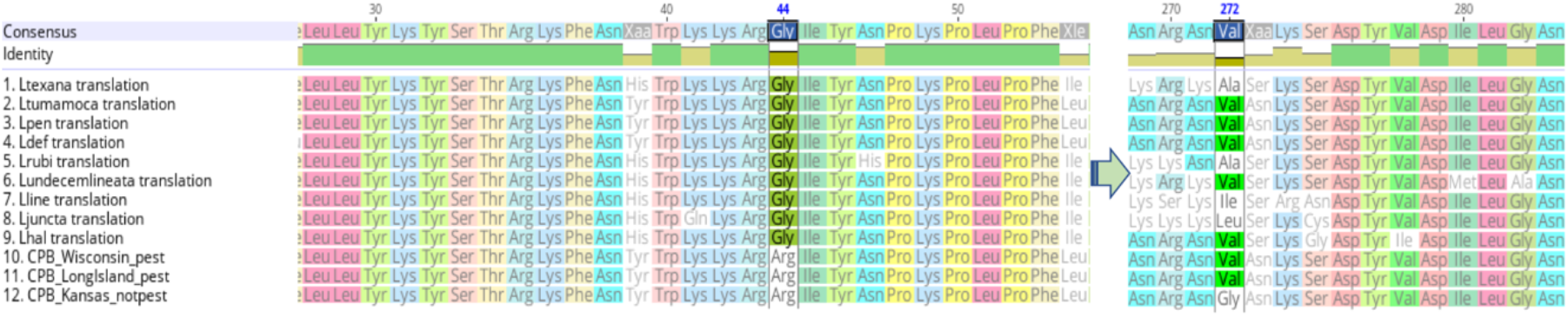
The shared significant cytochrome P450 locus, LDEC011604, with significant codon difference at position 44 and insignificant but shared in resistant CPB individuals (rows 10 and 11) at position 272.

### Testing for an Association between Genomic Features and Pestiferousness

We found that the phylogeny accounted for the total number of positively selected loci unique to each lineage, Blomberg’s K=1.45 (**Supplemental Table S6**, **Figure 5**), as well as a significant local level of autocorrelation (**Figure 2**). Furthermore, this node showed greater rates of trait evolution for total loci under selection (from the non-pest CPB individual) and resistant loci under selection (from the shared pest CPB individuals) after standardizing by genetic distance (**Supplemental Table S7**, **Supplemental Figure S8**). Additionally, the average percentage of candidate resistant loci under selection in the two pest CPB individuals was correlated with their average heterozygosity (p-value = 0.005506).

**Figure 5.**
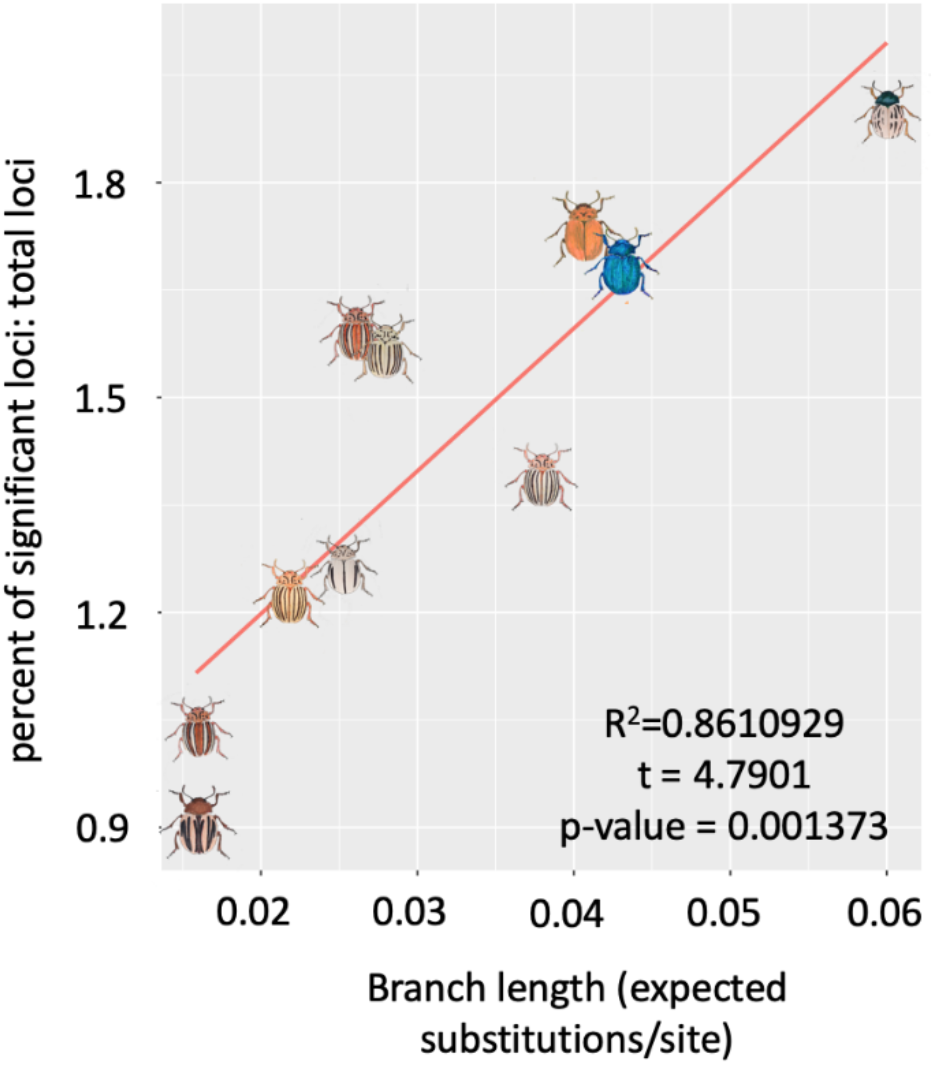
Proportion of loci under positive selection (results from adaptive branch-site random effect model) to total loci explained by branch length with significant phylogenetic dependence and accelerated rate of evolution.

## Discussion

### Genomic Features and Evolutionary Mechanisms of Pestiferousness

An outstanding question in pest evolution theory is how insects rapidly adapt to intense insecticidal pressure, new ecosystems and novel host plants. Previous comparative genomic research in model and non-model systems, including insect disease vectors, have associated different genomic feature with rapid adaptation. Empirical studies have shown the significance of gene family diversification (Hahn et al. 2007; Thomas et al. 2018; Gloss et al. 2019), protein-sequence evolution (Roux et al. 2014; Neafsey et al. 2015; Cicconardi et al. 2017) including convergent evolution (Chikina et al. 2016), and transposable element activity (Stapley et al. 2015) as important features. Here we employed comparative genomic techniques to test the significance of these mechanisms, as well as standing genetic diversity (Schoville et al. 2018), in order to describe evolutionary drivers of pestiferousness and rapid adaptation in an agriculturally impactful pest. We found little support for the role of gene family evolution, or for transposable element activity, driving evolutionary differences in the CPB pest lineage. We did, however, find support for both standing genetic diversity and rates of positive selection on protein coding genes.

### Unsupported mechanisms of rapid evolution

The determination of TE abundance was hypothesized to contribute to CPB adaptation (Schoville et al. 2018), however there was no indication that the levels of TE groups identified varied significantly among *Leptinotarsa* species. Only two TE families (tRNA and Gypsy-Cigr) changed in size significantly in our study, and these do not seem to be associated with rapid adaptation as they are simultaneously expanding and contracting in different lineages of *Leptinotarsa* (**Supplemental Table S3**).

Gene family size variation was considered as another driver for rapid adaptation. We anticipated large expansions in orthogroups clearly associated with host plant feeding and xenobiotic detoxification. While we did not find strong evidence for gene family turnover in *Leptinotarsa*, the significant orthogroup variation that we did observe shows functional overlap with our results from tests for positively selected genes (**Supplemental Table S1**). We found evidence that cysteine-type peptidase activity (GO:0008234) is a contracting orthogroup in CPB, despite previous work suggesting that these genes play a role in plant digestion in *L. decemlineata* (Schoville et al. 2018). This pattern and other notable gene family contractions might be linked to host-plant switches, as the two species with the most EOGs contracting, *L. peninsularis* and *L. tumamoca*, are distinctive in their preference for non-*Solanum* plants, including Wright’s groundcherry (*Physalis wrightii*) and Mexican poppy (*Kallstroemia grandiflora*), respectively (Jacques 1988).

### The importance of standing genetic variation in rapid adaptation

In order to test if standing genetic variation could be attributed to CPB success, we measured whether heterozygosity values were larger in ancestral, non-pest CPB individuals. Values in pest CPB populations might be biased as a consequence of founder events during range expansion and intense selection from various agricultural pressures. Indeed, our three CPB individuals show a predictive decrease in heterozygosity correlated with pest status, agroecosystem establishment, and spatial distance from center of origin (Southwestern US and Great Plains; Izzo et al. 2018). The non-pest CPB individual has the greatest heterozygosity, suggestive of a large effective population and high allelic diversity prior to agroecosystem invasion. One non-pest species, *L. haldemani*, also has relatively high heterozygosity. Notably, it is the only other *Leptinotarsa* species known to host on >3 genera of plants, and suggests that heterozygosity might be an important feature for host expansion or a consequence of greater niche breadth (Kassen 2002). Levels of heterozygosity, or standing genetic variation, have been linked to rapid adaptive evolution in a variety of study systems (Messer and Petrov 2013), including other cases of repeatedly evolving to toxic environments (Whitehead et al. 2017; Wilson et al. 2016). Importantly, it may be directly linked to our second mechanism of rapid adaptation, the rate of positive selection on protein-coding genes, as populations with high standing variation typically have higher rates of adaptive evolution (Barrett and Schluter 2008; Hermisson et al., 2008).

### Rates of positive selection in rapid adaptation

The number of genes under positive selection in the CPB lineage is significantly greater than what is expected based on its phylogenetic position. Although the cause of this elevated rate in CPB is not yet determined, it could be linked to high standing genetic diversity and/or larger effective population sizes. However, a high rate is not seen in the other polyphagous species, *L. haldemani*, despite large heterozygosity in this species. The high rates of positively selected genes observed in *L. tumamoca* and *L. peninsularis* are a bit puzzling but could be a result of intense selection associated with novel host plant expansion, as mentioned previously. A broader phylogenomic sampling of *Leptinotarsa*, or other species that vary in host use (e.g. Techer et al. 2019), needs to be studied in order to disentangle the relative importance of host switches on rates of positive selection. Furthermore, there is a clear pattern of selection on molecular pathways related to detoxification within the pest CPB lineages.

Recent experiments have shown dramatic regulatory changes in detoxification pathways following host shifts in plant-feeding arthropods (Dermauw et al., 2013; Zhang et al. 2017; Yu et al., 2016; Müller et al., 2017; Govind et al., 2010). Our analysis suggests that host shifts eventually lead to protein-coding changes during the process of speciation. The enriched GO terms leading to the Colorado potato beetle clade (**Table 3**, nodes 5-8), includes nuclease activity, which includes enzymes for plant digestion and detoxification of plant secondary metabolites, and general molecular binding, which plays a role in rapid regulatory evolution (Schernthaner et al. 2002). Other comparative genomic studies have found similar evidence of positive selection in detoxification genes related to hostplant utilization (Simon et al. 2015), but have typically involved comparisons of highly divergent taxa (but see Gloss et al., 2019, as an exception). While positive selection is evident throughout the phylogeny, enrichment of molecular functions related to pest traits are only evident in pest CPB genomes. Significant genes in the branch-site test for positive selection are clearly associated with agricultural adaptation and pestiferousness.

Within pest CPB individuals, there is enrichment of several interesting molecular functions that could be related to agricultural adaptation (**Table 4**). First, superoxide dismutase (SOD) activity (GO:0004784 and 0019430), an enzyme that functionally mitigates oxidative stress at the cellular level, could play an important role in CPB adaptation to environmental and chemical stress (Abdollahi et al. 2004; Velki et al. 2011). In addition to mitigating effects of insecticide exposure, SOD may be co-opted by diapausing beetles to persist throughout the winter as an additional protective measure for longevity (Sim and Denlinger 2011). Second, the expansive group of membrane bound proteins (GO:0016021) has many genes associated with pestiferousness such as: hormone receptors and transporters, chemosensory receptors, and detoxification enzymes (**Supplemental Table S5**). Behaviors of foraging, copulation, and aggression are important phenotypes associated with hormone physiology (Ma, Guo and Kang, 2016; Burggren, 2017, Braendle and Flatt, 2006). The selective pressure of environmental stress or intraspecific competition in novel agroecosystems might have driven the observed variation at octopamine receptor (LDEC006841-RA) and dopamine transporter (LDEC004177-RA) in the pest CPB (**Supplemental Table S5)** (van Straalen et al. 2004). Chemosensory adaptation involving positive selection of several gustatory receptors in the pest-CPB individuals, is also intriguing. Gustatory receptors stimulate feeding by binding to nutrients in leaf tissue and have been well described in the Colorado potato beetle system (Ishikawa et al., 1969; Mitchell 1985; Mitchell and McCashin 1994; Mitchell and Schoonhoven 1974). The physiology of the pest-specific gustatory receptors is yet to be determined and could be an interesting avenue in exploring preference and performance of CPB on potato.

We further analyzed the candidate resistant loci, under positive selection, between pest and non-pest CPB. Although, the GO terms associated with these enzymes and receptors were not enriched in non-pest CPB, as they were in both pest-CPB individuals, there were still several genes under selection in the non-pest individual. All three samples share four candidate resistant genes, while the two pest CPB share an additional two candidate resistant genes (**Table 5**). The first significant resistant gene shared by all three CPB’s is a 305 alpha-1 like cytochrome P450 (LDEC016278) and is likely associated with the metabolism of hormones, not xenobiotics, as it is highly expressed in the spermathecae of *Drosophila melanogaster* (Niwa et al. 2004). Of the four shared and significant loci, the second discussed here is the ryanodine calcium channel LDEC018687. Ryanodine calcium channels are target sites for both botanical insecticides and the naturally occurring alkaloid ryanodine (Wan et al. 2014). Interestingly, target-site knockdown of these receptors has already been associated with resistance to ryanodine insecticide formulations in the Colorado potato beetle (Wan et al., 2014), but the shared significance for this locus among pest and non-pest CPB would most likely reflect ancestral phyto-ryanodine exposure. The shared glutamate receptor, LDEC007559, could also be a consequence of natural plant defensive chemistry exposure, and the venom carboxylesterase, LDEC005267, is annotated as an allergen expressed in the venom duct of bees. This ortholog may or may not be a serine esterase that has been coopted for defense by the CPB, as it has in *Apis mellifera* (Elsik et al. 2014). In summary, the shared loci resulting from positive selection in the CPB lineage do not suggest pre-adaptive mechanisms for the rapid evolution observed in pest-CPB populations when exposed to insecticides.

However, the two loci uniquely under selection in the pest CPB individuals, an esterase (LDEC014122) and a cytochrome P450 (LDEC011604), are most likely the result of positive selection in an agricultural setting. The esterase has been associated with insecticide detoxification of an organophosphate insecticide, a common legacy agricultural control agent, in *Locusta migratoria* (Zhang et al. 2013). At this gene, the pest and non-pest CPB share two significant sites that vary: codon 14 as phenylalanine → threonine and codon 18 as leucine → valine (**Figure 3**); additionally, the pest CPB share another significant codon change at position 16, valine → methionine, which could be due to recent common-ancestry or independent and convergent selection from organophosphate insecticide exposure. The pest-specific CYP p450, LDEC011604, has also been linked with neonicotinoid resistance in CPB and other insect pests (Clements et al., 2018). Upregulation of CYP 6k1 locus has previously been observed in resistant Colorado potato beetles from Wisconsin when exposed to both the insecticide imidacloprid, a neonicotinoid, and the broad-spectrum fungicide chlorothalonil (Clements et al. 2018). At the codon level, there were shared significant changes among the three CPB individuals, codon 44 glycine → arginine, and one insignificant, yet shared codon changes among the pests at position 272 (**Figure 4**).

### Additional Insights from Comparative Genomics of Leptinotarsa

The divergence events in the genus *Leptinotarsa* contribute to our understanding of how plant-feeding insects evolve, particularly whether speciation is associated with environmental change or host-plant diversification (Mitter et al. 1988; Kergoat et al. 2017). Our time calibrated *Leptinotarsa* phylogeny suggests that divergence of the common ancestor occurred around 19 Ma, with the clade including the Colorado potato beetle arising ~10 Ma. Intriguingly, this period includes notable diversification events within the Solanaceae, such as the split of *Solanum* & *Nicotiana* at ~24 Ma, *Solanum* & *Capsicum* at ~19 Ma, and tomato, *S. lycopersicum*, and potato, *L. tuberosum*, at ~8 Ma (Särkinen et al. 2013). As *Leptinotarsa* species are mostly monophagous on Solanaceae, this pattern might suggest co-diversification with *Solanum* host plants. However, as host generalization appears to have evolved multiple times in our phylogenetic reconstruction, it seems unlikely that co-speciation with host plants is the main mechanism driving diversification in *Leptinotarsa*, and rather, these beetles most likely radiate onto existing hostplant diversity (Gómez-Zurita et al. 2007). Alternative drivers of speciation include climatic events during the Miocene epoch (~23.03-5.33 Ma), when global temperatures became significantly warmer and drier, and geological events associated with the orogeny of the Sierra Madre de Chiapas in southern Mexico ~12-10 Ma, possibly driving speciation via allopatric isolation (Witt et al. 2012). Additional sampling of *Leptinotarsa* and related taxa, as well as population-level sampling, will be required to test these competing hypotheses regarding plant-insect co-speciation, climate change, and geology.

## Conclusion

We employed phylogenomic comparisons to test for mechanisms underlying the rapid evolution of pestiferousness within the Colorado potato beetle lineage. Our results suggest that high heterozygosity and elevated rates of positive selection on protein coding genes, particularly genes associated with agroecosystem adaptation (insecticide resistance and xenobiotic metabolism, chemosensation, and hormone pathways) appear to be unique features of the Colorado potato beetle. Furthermore, the genetic diversity and directional selection characteristic of the *Leptinotarsa* genus may have facilitated the evolution of pestiferousness in the Colorado potato beetle. Performing similar phylogenetic analyses on other notable pest species, and their relatives, should add valuable context to the idea that novel genomic features can predispose a species to pestiferousness.

## Acknowledgements

The authors wish to thank Margarethe Brummerman for help in obtaining samples of *Leptinotarsa*. Helpful comments on the manuscript were provided by Cécile Ané, Carol E. Lee, Russell Groves, Lyric Bartholomay, David Baum, Michael S. Crossley, Justin Clements, Ben Bradford, as well as Joshua, Alisa and Richard Cohen. This research was funded by a USDA NIFA Exploratory grant (2015-67030-23495) and a USDA Hatch grant (WIS02004).

## Data Accessibility

Genomic data are publicly available at NCBI on Genbank (Bioproject: PRJNA580490), as raw data in the short read archive (accessions SRR10388401:SRR10388312) and whole genome shotgun accessions (WOGI00000000:WOGR00000000).

