## Supplemental Materials for "Elevated rates of positive selection drive the evolution of pestiferousness in the Colorado potato beetle (*Leptinotarsa decemlineata*, Say)"

**Supplemental Table S1.** Significant gene families experiencing copy number variation at the species level. Bolded GO terms correspond to those enriched from Fisher-exact test and aBSREL results as well as those annotated in orthogroups. Green for unique GO terms in expanding orthogroups, yellow for GO terms in orthogroups expanding/contracting in multiple lineages and red for GO terms in uniquely contracting orthogroups.

| *species* | *OGs expanding* | *GO term annotations* | *OGs contracting* | *GO term annotations* |
|---|---|---|---|---|
| *L. texana* | EOG8M945N[+5] | GO:0046872: metal ion binding | EOG85XB8W [-19] | **GO:0003777: microtubule motor activity** |
| | | **GO:0003676: nucleic acid binding** | | **GO:0005524: ATP binding** |
| | | | | GO:0007018: microtubule-based movement |
| | | | | GO:0030286: dynein complex |
| | EOG8KPWS1[+5] | GO:0003723: RNA binding | EOG8577MX [-6] | GO:0046872: metal ion binding |
| | | GO:0008033: tRNA processing | | |
| | | GO:0008168: methyltransferase activity | | GO:0006486: protein glycosylation |
| | | GO:0006364: rRNA processing | | GO:0005794: Golgi apparatus |
| | | | | GO:0016757: transferase activity, transferring glycosyl groups |
| | | | | **GO:0016021: integral component of membrane** |

| | | | | |
|---|---|---|---|---|
| *L. juncta* | EOG8WDGTK[+9] | GO:0005198: structural molecule activity | EOG8HDWBC [-8] | MADF domain |
| | | GO:0019028: viral capsid | | |
| | EOG8RZ1FV [+7] | **GO:0005524: ATP binding** | | |
| | EOG8VQD39 [+6] | GO:0046872: metal ion binding | | |
| | | **GO:0003676: nucleic acid binding** | | |
| *L. undecemlineata* | | | EOG82Z74B [-7] | **GO:0005524: ATP binding** GO:0046872: metal ion binding |
| | | | | GO:0004674: protein serine/threonine kinase activity |
| | | | | GO:0035556: intracellular signal transduction |
| | | | | GO:0046872: metal ion binding |
| *L. defecta* | | | EOG86T5FN [-5] | **GO:0003676: nucleic acid binding** |
| | | | | GO:0046872: metal ion binding |
| *L. decemlineata* | | | EOG8P2SH4 [-7] | GO:0008234: cystei`ne-type peptidase activity |
| *L. peninsularis* | EOG86T5FN [+6] | **GO:0003676: nucleic acid binding** | EOG8Z91BC [-12] | GO:0046872: metal ion binding |
| | | GO:0046872: metal ion binding | | **GO:0003676: nucleic acid binding** |
| | EOG8J9QCN [+4] | **GO:0005524: ATP binding** | EOG8Z91BR [-14] | **GO:0005524: ATP binding** |
| | | **GO:0016887: ATPase activity** | | **GO:0003700: sequence-specific DNA** |

| | | | | |
|---|---|---|---|---|
| | | | | **binding transcription factor activity** |
| | | | | **GO:0003677: DNA binding** |
| | | | | GO:0006351: transcription, DNA-templated |
| | | | | **GO:0005634: nucleus** |
| | | | EOG8Z919V [-14] | GO:0008241: peptidyl-dipeptidase activity |
| | | | | GO:0008237: metallopeptidase activity |
| | | | | GO:0016020: membrane |
| | | | EOG851H68 [-7] | IPR012337: Ribonuclease H-like domain IPR018289: MULE transposase domain |
| | | | EOG8Z91BS [-9] | IPR006600: HTH CenpB-type DNA-binding domain IPR004875: DDE superfamily endonuclease, CENP-B-like |
| | | | EOG8Z91D3 [-6] | **GO:0003676: nucleic acid binding** |
| | | | | **GO:0008270: zinc ion binding** |
| | | | EOG8Z91BV [-7] | IPR010512: Protein of unknown function DUF1091 |

| | | | | |
|---|---|---|---|---|
| | | | EOG8Z919Z [-5] | GO:0046873: metal ion transmembrane transporter activity |
| | | | | GO:0016020: membrane |
| | | | EOG8Z91CV [-8] | **GO:0043565: sequence-specific DNA binding** |
| | | | | **GO:0003700: sequence-specific DNA binding transcription factor activity** |
| | | | | **GO:0005634: nucleus** |
| | | | EOG8Z919F [-4] | -- |
| | | | EOG8Z91B7 [-7] | GO:0005887: integral component of plasma membrane |
| | | | EOG8Z0DR4 [-6] | -- |
| | | | EOG8Z28XG [-3] | -- |
| *L. tumamoca* | EOG8QVFSS [+4] | **GO:0000166: nucleotide binding** | EOG8VQD39 [-5] | GO:0046872: metal ion binding |
| | | | | **GO:0003676: nucleic acid binding** |
| | | **GO:0003676: nucleic acid binding** | EOG8Q2GVQ [-10] | GO:0046872: metal ion binding |
| | | | | **GO:0003676: nucleic acid binding** |
| | EOG84J4XX [+7] | **GO:0003777: microtubule motor activity** | EOG8QC3KD [-6] | GO:0005543: phospholipid binding |
| | | | | GO:0005737: cytoplasm |
| | | **GO:0005524: ATP binding** | EOG8Q2GWN [-7] | GO:0003723: RNA binding |

| | | | | |
|---|---|---|---|---|
| | | | | **GO:0006396: RNA processing** |
| | | GO:0007018: microtubule-based movement | EOG8Q2GW6 [-5] | GO:0003995: acyl-CoA dehydrogenase activity |
| | | | | GO:0050660: flavin adenine dinucleotide binding |
| | | GO:0005871: kinesin complex | EOG8Q2GVS [-5] | GO:0046872: metal ion binding |
| | | | | **GO:0003677: DNA binding** |
| | | GO:0005874: microtubule | EOG8QC3KJ [-5] | IPR001478: PDZ domain<br>IPR001849: Pleckstrin homology domain |
| *L. rubiginosa* | EOG8TQPPS [+6] | GO:0005086: ARF guanyl-nucleotide exchange factor activity | EOG8WDGTK [-9] | GO:0005198: structural molecule activity |
| | | GO:0032012: regulation of ARF protein signal transduction | | GO:0019028: viral capsid |
| | EOG8KPWS1[+6] | GO:0003723: RNA binding | | |
| | | GO:0008168: methyltransferase activity | | |
| | | GO:0006364: rRNA processing | | |
| | | GO:0008033: tRNA processing | | |
| *L. haldemani* | EOG8M945N [+6] | GO:0046872: metal ion binding | EOG80GG63 [-6] | |
| | | **GO:0003676: nucleic acid binding** | | |

| | | | |
|---|---|---|---|
| | | EOG80GG4N [-9] | GO:0005254: chloride channel activity |
| | | | GO:0034707: chloride channel complex |
| | | EOG8TQPPS [-6] | GO:0005086: ARF guanyl-nucleotide exchange factor activity |
| | | | GO:0032012: regulation of ARF protein signal transduction |
| | | EOG80GG58 [-7] | GO:0046872: metal ion binding |
| | | | **GO:0003676: nucleic acid binding** |
| *L. lineolata* | n/a | n/a | |

**Supplemental Table S2.** All annotated TE elements, with percent abundance of genome, among ten *Leptinotarsa* beetles compared.

| **Transposable Elements in the Genomes of *Leptinotarsa* species (percent of genome)** | | | | | | | | | |
|---|---|---|---|---|---|---|---|---|---|
| **TE Class** | ***L.defecta*** | ***L.haldemani*** | ***L.juncta*** | ***L.lineolata*** | ***L.peninsularis*** | ***L.rubiginosa*** | ***L.texana*** | ***L.tumamoca*** | ***L.undecemlineata*** |
| Academ | 0.59 | 0.55 | 0.36 | 0.42 | 0.37 | 0.37 | 0.3 | 0.43 | 0.43 |
| CMC-Chapaev-3 | 0.28 | 0.39 | 0.31 | 0.68 | 0.21 | 0.26 | 0.3 | 0.22 | 0.31 |
| CMC-EnSpm | 0.13 | 0.07 | 0.01 | 0.04 | 0.07 | 0.01 | 0.03 | 0.01 | 0.01 |
| CMC-Transib | 0.01 | 0.01 | 0.01 | 0 | 0 | 0.01 | 0.05 | 0 | 0 |
| Copia | 0.1 | 0.07 | 0.05 | 0.1 | 0.11 | 0.09 | 0.09 | 0.07 | 0.32 |
| CR1 | 1.16 | 1.21 | 0.94 | 1.13 | 0.89 | 0.91 | 1.3 | 1.78 | 1.11 |
| CR1-Zenon | 2.18 | 1.33 | 1.88 | 2.59 | 2.19 | 1.97 | 1.79 | 1.32 | 1.94 |
| CRE | 0.12 | 0.04 | 0 | 0 | 0 | 0 | 0.03 | 0.07 | 0 |
| CRE-II | 0.1 | 0.29 | 0.15 | 0.16 | 0.11 | 0.19 | 0.2 | 0.08 | 0.15 |
| Crypton | 0.47 | 0.35 | 0.35 | 0.69 | 0.38 | 0.39 | 0.42 | 0.4 | 0.44 |
| Crypton-V | 0 | 0 | 0 | 0 | 0 | 0 | 0 | 0.01 | 0 |
| Dada | 0 | 0 | 0 | 0 | 0 | 0 | 0 | 0 | 0.01 |
| DIRS | 0.01 | 0 | 0 | 0 | 0.01 | 0.01 | 0.03 | 0.01 | 0.01 |
| DNA | 0.84 | 0.46 | 0.69 | 0.71 | 0.47 | 0.65 | 0.53 | 0.21 | 0.89 |
| Dong-R4 | 0.96 | 1.08 | 1.03 | 1.22 | 0.78 | 0.67 | 0.97 | 1 | 1.35 |
| ERV1 | 0 | 0 | 0.02 | 0.01 | 0 | 0 | 0 | 0.01 | 0 |
| Ginger | 0.01 | 0.01 | 0.01 | 0 | 0 | 0 | 0.06 | 0.01 | 0 |
| Gypsy | 0.47 | 0.37 | 0.46 | 0.86 | 1.28 | 0.39 | 0.47 | 0.44 | 0.54 |
| Gypsy-Cigr | 0.21 | 0.06 | 0 | 0.12 | 0.18 | 0.07 | 0.05 | 0.09 | 0.2 |
| hAT | 0 | 0 | 0 | 0 | 0.06 | 0 | 0 | 0 | 0 |
| hAT-Ac | 0.08 | 0.06 | 0.03 | 0.11 | 0.08 | 0.04 | 0.06 | 0.06 | 0.06 |
| hAT-Blackjack | 0 | 0.02 | 0.02 | 0.02 | 0 | 0.01 | 0 | 0.01 | 0.01 |
| hAT-Charlie | 0.43 | 0.42 | 0.44 | 0.47 | 0.3 | 0.34 | 0.67 | 0.35 | 0.49 |
| hAT-hATm | 0.03 | 0.03 | 0.04 | 0.03 | 0.05 | 0.08 | 0.02 | 0.03 | 0 |
| hAT-Tag1 | 0 | 0 | 0 | 0 | 0 | 0 | 0 | 0.01 | 0.01 |
| hAT-Tip100 | 0.23 | 0.25 | 0.15 | 0.19 | 0.13 | 0.13 | 0.18 | 0.15 | 0.31 |
| Helitron | 0.32 | 0.37 | 0.32 | 0.59 | 0.36 | 0.35 | 0.35 | 0.31 | 0.33 |
| I | 0.19 | 0.19 | 0.13 | 0.14 | 0.12 | 0.16 | 0.19 | 0.12 | 0.14 |

| | | | | | | | | | |
|---|---|---|---|---|---|---|---|---|---|
| I-Nimb | 0 | 0.01 | 0.04 | 0.11 | 0.01 | 0.02 | 0.02 | 0.01 | 0.06 |
| Jockey | 0.59 | 0.85 | 0.57 | 0.92 | 0.49 | 0.6 | 0.73 | 0.54 | 0.63 |
| Kolobok | 0 | 0.01 | 0 | 0 | 0 | 0 | 0 | 0 | 0 |
| Kolobok-Hydra | 0 | 0 | 0.01 | 0 | 0 | 0 | 0 | 0 | 0 |
| L1 | 0 | 0 | 0 | 0.01 | 0.09 | 0 | 0.03 | 0 | 0 |
| L1-Tx1 | 0.01 | 0.01 | 0.01 | 0 | 0 | 0.01 | 0.03 | 0.01 | 0 |
| L2 | 4.13 | 4.2 | 3.67 | 4.84 | 3.41 | 3.39 | 4.84 | 3.51 | 4.9 |
| LINE | 0.04 | 0.04 | 0.03 | 0.14 | 0.13 | 0.09 | 0.06 | 0.15 | 0.06 |
| LOA | 3.31 | 3.14 | 2.96 | 1.78 | 3.29 | 2.74 | 3.22 | 3.18 | 3.58 |
| LTR | 0.04 | 0.04 | 0.01 | 0 | 0.01 | 0.02 | 0.05 | 0 | 0.01 |
| Maverick | 0.65 | 0.15 | 0.34 | 0.53 | 0.47 | 0.41 | 1.03 | 0.4 | 0.53 |
| Merlin | 0.01 | 0.01 | 0 | 0 | 0 | 0 | 0.01 | 0 | 0 |
| MULE-MuDR | 0 | 0 | 0.01 | 0 | 0 | 0 | 0 | 0 | 0.02 |
| MuLE-NOF | 0.01 | 0 | 0 | 0 | 0 | 0.01 | 0 | 0 | 0 |
| Ngaro | 0 | 0.01 | 0 | 0.02 | 0 | 0 | 0.02 | 0 | 0 |
| P | 0.1 | 0.09 | 0.07 | 0.27 | 0.09 | 0.07 | 0.11 | 0.06 | 0.08 |
| Pao | 0.23 | 0.16 | 0.17 | 0.19 | 0.21 | 0.14 | 0.41 | 0.16 | 0.2 |
| Penelope | 4.86 | 4.91 | 4.52 | 4.01 | 4.87 | 5.18 | 5.76 | 5.28 | 4.78 |
| PIF-Harbinger | 0.1 | 0.17 | 0.52 | 0.03 | 0.09 | 0.51 | 0.62 | 0.43 | 0.1 |
| PIF-ISL2EU | 0.05 | 0.04 | 0.01 | 0.06 | 0 | 0.03 | 0.06 | 0 | 0 |
| PiggyBac | 0.03 | 0.07 | 0.04 | 0.11 | 0 | 0.03 | 0.07 | 0.04 | 0.14 |
| R1 | 0.16 | 0.21 | 0.15 | 0.2 | 0.15 | 0.13 | 0.16 | 0.12 | 0.16 |
| R2 | 0.01 | 0 | 0 | 0.01 | 0.01 | 0.01 | 0 | 0 | 0 |
| Rex-Babar | 0 | 0 | 0 | 0 | 0.01 | 0 | 0 | 0.01 | 0.02 |
| RTE-BovB | 0.04 | 0.02 | 0.04 | 0.02 | 0.01 | 0.01 | 0.06 | 0.01 | 0.03 |
| RTE-X | 0 | 0 | 0 | 0.01 | 0 | 0 | 0.01 | 0.01 | 0.01 |
| SINE | 0.01 | 0 | 0.01 | 0.25 | 0.01 | 0 | 0.01 | 0.01 | 0.01 |
| Sola | 0.07 | 0.01 | 0.04 | 0.07 | 0.06 | 0.02 | 0.01 | 0 | 0.02 |
| Tad1 | 0 | 0 | 0 | 0.1 | 0 | 0.18 | 0.1 | 0.03 | 0 |
| TcMar | 0.02 | 0.01 | 0 | 0 | 0 | 0 | 0.01 | 0 | 0 |
| TcMar-Fot1 | 0.05 | 0.07 | 0.06 | 0.1 | 0.03 | 0.04 | 0.21 | 0.05 | 0.1 |
| TcMar-m44 | 1.03 | 0.77 | 0.75 | 0.9 | 0.83 | 0.59 | 0.94 | 0.57 | 0.83 |
| TcMar-Mariner | 2.67 | 1.75 | 2.32 | 2.62 | 2.7 | 3.18 | 2.04 | 2.67 | 2.39 |
| TcMar-Pogo | 0 | 0 | 0 | 0 | 0.01 | 0 | 0 | 0.01 | 0.01 |
| TcMar-Sagan | 0 | 0 | 0 | 0 | 0 | 0 | 0.01 | 0 | 0 |
| TcMar-Tc1 | 4.57 | 3.7 | 3.27 | 6.16 | 3.6 | 3.55 | 4.69 | 4.04 | 5.22 |
| TcMar-Tigger | 0.02 | 0.01 | 0 | 0 | 0 | 0.01 | 0 | 0 | 0 |
| tRNA | 0.18 | 0.25 | 0.05 | 0.05 | 0.06 | 0.03 | 0.08 | 0.25 | 0 |
| Zator | 0.03 | 0.03 | 0.03 | 0.01 | 0.03 | 0.03 | 0.03 | 0.03 | 0.02 |

| Total | 31.94 | 28.37 | 27.1 | 33.8 | 28.82 | 28.13 | 33.52 | 28.78 | 32.97 |
|---|---|---|---|---|---|---|---|---|---|

**Supplemental Table S3**. TE annotations experiencing rapid copy number variation in *Leptinotarsa*.

| *species* | *TEs expanding* | *TEs contracting* |
|---|---|---|
| *L. decemlineata* | Gypsy-Cigr [+5] | tRNA [-8] |
| *L. juncta* | -- | Gypsy-Cigr [-6] |
| *L. haldemani* | tRNA [+18] | -- |
| *L. tumamoca* | tRNA [+16] | -- |

**Supplemental Table S4**. Positively selected genes shared among all three CPB conspecifics genes with enriched GO terms.

| | |
|---|---|
| LDEC000730-RA | lon protease mitochondrial isoform x2\|GO:0004176 GO:0004252 GO:0004729 GO:0005524 GO:0006515 GO:0006779 GO:0055114 GO:0006510 GO:0015994 |
| LDEC002335-RA | trna (uracil-5-)-methyltransferase homolog a-like isoform x1\|GO:0000166 GO:0003676 GO:0008173 GO:0001510 GO:0006396 |
| LDEC004614-RA | ubiquitin carboxyl-terminal hydrolase 31-like\|GO:0036459 GO:0006511 GO:0016579 |
| LDEC004994-RA | endothelial differentiation-related factor 1 homolog\|GO:0005634 GO:0005737 GO:0003713 GO:0008327 GO:0030674 GO:0007417 GO:0007424 GO:0045944 GO:0048813 GO:0005667 GO:0045941 |
| LDEC007808-RA | magnesium transporter NIPA2\|GO:0016021 GO:0015095 GO:0015693 GO:1903830 |
| LDEC008531-RA | 26s proteasome non-atpase regulatory subunit 4\|GO:0008540 GO:0006511 |
| LDEC009099-RA | geranylgeranyl pyrophosphate synthase\|GO:0016021 GO:0004311 GO:0006694 GO:0016114 |
| LDEC012472-RA | aprataxin\|GO:0003677 GO:0005515 GO:0033699 GO:0006281 |
| LDEC016151-RA | 3-hydroxy-3-methylglutaryl- reductase\|GO:0005789 GO:0016021 GO:0004420 GO:0050661 GO:0008299 GO:0015936 GO:0055114 GO:0006694 |
| LDEC018296-RA | probable low-specificity l-threonine aldolase 2\|GO:0004793 GO:0006544 GO:0006563 GO:0006566 |
| LDEC021484-RA | chondroitin sulfate N-acetylgalactosaminyltransferase 1\|GO:0016021 GO:0032580 GO:0008376 GO:0008152 |
| LDEC021876-RA | probable E3 ubiquitin- ligase sinah\|GO:0005634 GO:0005515 GO:0008270 GO:0006511 GO:0007275 |
| LDEC023421-RA | protocadherin-like wing polarity protein stan isoform x1\|GO:0005886 GO:0005911 GO:0016021 GO:0004930 GO:0005057 GO:0005509 GO:0007156 GO:0007186 GO:0007367 GO:0016318 GO:0016319 GO:0016339 GO:0019233 GO:0030111 GO:0035159 GO:0045746 GO:0045773 GO:0048057 GO:0048813 GO:0051963 GO:0070593 GO:0090175 GO:1902669 |

**Supplemental Table S5**. Positively selected genes in pest CPB genes with enriched GO terms.

| | |
|---|---|
| LDEC000240-RA | facilitated trehalose transporter tret1 isoform x1\|GO:0016021 GO:0022891 GO:0055085 |
| LDEC000473-RA | flocculation protein flo11\|GO:0016021 |
| LDEC001548-RA | UNC93 isoform X1\|GO:0016021 |
| LDEC001619-RA | arginine N-methyltransferase 7\|GO:0008168 GO:0006479 |
| LDEC001672-RA | mucin-17 isoform X1\|GO:0016021 GO:0004725 GO:0035335 GO:0006570 |

| | |
|---|---|
| LDEC001696-RA | 5-hydroxytryptamine (serotonin) receptor\|GO:0016021 GO:0004930 GO:0007186 |
| LDEC002144-RA | atp-binding cassette sub-family b member mitochondrial-like\|GO:0016021 GO:0005524 GO:0042626 GO:0008152 GO:0055085 |
| LDEC002874-RA | protein arginine n-methyltransferase 1\|GO:0008168 GO:0006479 |
| LDEC002928-RA | tmem9 family protein\|GO:0016021 |
| LDEC003420-RA | junctophilin-1 isoform x1\|GO:0016021 GO:0006810 |
| LDEC004061-RA | sex determination protein fruitless isoform x1\|GO:0005634 GO:0003677 GO:0003700 GO:0046872 GO:0002118 GO:0007417 GO:0007517 GO:0007530 GO:0007620 GO:0016199 GO:0016543 GO:0044719 GO:0045433 GO:0046661 GO:0048047 GO:0048813 GO:0005667 GO:0045449 |
| LDEC004177-RA | high-affinity dopamine transporter partial\|GO:0016021 GO:0005328 GO:0006836 GO:0055085 GO:0006812 |
| LDEC004625-RA | agap010471-pa-like protein\|GO:0016021 |
| LDEC004879-RB | gustatory receptor r6\|GO:0005886 GO:0016021 GO:0050909 |
| LDEC004898-RA | glucosylceramidase-like\|GO:0004348 GO:0005975 GO:0006687 |
| LDEC005020-RA | cuticular protein analogous to peritrophins 1-b\|GO:0005576 GO:0016021 GO:0008061 GO:0006030 |
| LDEC005212-RA | neuroligin- partial\|GO:0016021 |
| LDEC005581-RA | actin-related protein 2 3 complex subunit 2\|GO:0005840 GO:0005885 GO:0045179 GO:0070938 GO:0003735 GO:0003779 GO:0005200 GO:0006412 GO:0008360 GO:0030031 GO:0030866 GO:0034314 GO:0046331 GO:0042254 |
| LDEC006551-RA | Glycerate kinase\|GO:0044237 |
| LDEC006841-RA | Octopamine receptor beta-1R\|GO:0016021 GO:0004930 GO:0007186 |
| LDEC006953-RA | disulfide-isomerase partial\|GO:0016021 GO:0016853 GO:0008152 GO:0045454 |
| LDEC007317-RA | alpha-aminoadipic semialdehyde dehydrogenase\|GO:0005634 GO:0005759 GO:0005829 GO:0070062 GO:0004043 GO:0005515 GO:0008802 GO:0006081 GO:0006554 GO:0007605 GO:0019285 GO:0055114 GO:0009085 GO:0006544 GO:0006563 GO:0006566 |
| LDEC007707-RA | nicotinic acetylcholine receptor subunit alpha4\|GO:0016021 GO:0030054 GO:0045211 GO:0004889 GO:0098655 GO:0007165 |
| LDEC008491-RA | zinc transporter zip3\|GO:0016020 GO:0046873 GO:0030001 GO:0055085 |
| LDEC010877-RA | teneurin-m isoform x4\|GO:0016021 GO:0005515 GO:0097264 |
| LDEC010928-RA | membrane-associated progesterone receptor component 2\|GO:0016021 |
| LDEC010997-RA | formin-like protein cg32138 isoform x2\|GO:0003779 GO:0017048 GO:0030036 |
| LDEC012537-RA | metabotropic glutamate receptor 2 isoform x2\|GO:0016021 GO:0004930 GO:0007186 |
| LDEC013042-RA | dolichyl-diphosphooligosaccharide--protein glycosyltransferase subunit 1\|GO:0016021 GO:0004579 GO:0008250 GO:0018279 |
| LDEC013235-RA | superoxide dismutase\|GO:0005615 GO:0005737 GO:0016020 GO:0004784 GO:0005507 GO:0008270 GO:0019430 GO:0055114 |
| LDEC014092-RA | PREDICTED: uncharacterized protein LOC100742412\|GO:0005737 GO:0016021 GO:0019991 GO:0035151 |
| LDEC014122-RA | esterase\|GO:0016021 GO:0016787 GO:0008152 |
| LDEC014687-RA | lachesin isoform x1\|GO:0016021 GO:0005515 |
| LDEC015298-RA | alpha-2a adrenergic receptor\|GO:0016021 GO:0004930 GO:0007186 |
| LDEC016656-RA | nucleolar and coiled-body phosphoprotein 1 isoform x3\|GO:0031981 |
| LDEC016726-RA | pyrroline-5-carboxylate reductase\|GO:0016021 GO:0004735 GO:0055114 GO:0055129 GO:0006525 |

| | |
|---|---|
| LDEC017126-RA | glutamine synthetase 2 cytoplasmic isoform x1\|GO:0016021 GO:0004356 GO:0005328 GO:0005524 GO:0006542 GO:0006836 GO:0055085 GO:0009252 GO:0006812 |
| LDEC018425-RA | transmembrane and immunoglobulin domain-containing protein\|GO:0016021 GO:0005515 |
| LDEC018725-RA | gustatory receptor 28b\|GO:0016021 GO:0050909 |
| LDEC018725-RB | gustatory receptor isoform D [Drosophila melanogaster]\|GO:0016021 GO:0050909 |
| LDEC018725-RC | gustatory receptor 88 [Tribolium castaneum]\|GO:0016021 GO:0050909 |
| LDEC018727-RA | gustatory receptor 2a\|GO:0016021 GO:0050909; gustatory receptor isoform B [Drosophila melanogaster]\|; AAEL011073- partial [Aedes aegypti]\| |
| LDEC018949-RA | dolichyl pyrophosphate Man9 c2 alpha-1,3-glucosyltransferase\|GO:0005789 GO:0016758 GO:0016772 GO:0044237 |
| LDEC023090-RA | esterase\|GO:0016787 GO:0008152 |
| LDEC024733-RA | low-density lipoprotein receptor-related protein 2\|GO:0016021 GO:0005509 GO:0005515 |

**Supplemental Table S6.** Significance levels (*p*-values) of *Leptinotarsa* features and traits under various models of phylogenetic dependence and autocorrelation.

| **Genomic features** | **Cmean** | **Moran's I** | **Blomberg's K** | **Blomberg's K.star** | **Pagel's Lambda** |
|---|---|---|---|---|---|
| Candidate Resistant loci Proportion | 0.418 | 0.521 | 0.667 | 0.609 | 1.000 |
| Host Breadth | 0.275 | 0.487 | 0.527 | 0.469 | 1.000 |
| Heterozygosity | 0.084 | 0.056 | 0.242 | 0.098 | 0.001 |
| Total loci proportion | 0.007 | 0.004 | 0.005 | 0.002 | 0.001 |

**Supplemental Table S7.** Standardized phylogenetic independent contrast values for total percentage of loci under selection and candidate insecticide resistant loci under selection (outlier node in red, ancestral to *L. decemlineata, L. tumamoca and L. peninsularis*.

| Node number | Standardized total loci PIC (abs(PIC)/sqrt(Σ branch lengths)) | Standardized resistant loci PIC (abs(PIC)/sqrt(Σ branch lengths)) |
|---|---|---|
| 1 | 2.960252708 | 0.836517373 |
| 2 | 2.263019995 | 2.450955612 |
| 3 | 5.141949065 | 2.542008412 |
| 4 | 3.319603589 | 2.496642682 |
| 5 | 5.871454231 | 8.156845247 |
| 6 | 1.482570815 | 1.764777081 |
| 7 | 2.733607017 | 0 |
| 8 | 8.759990083 | 18.31869729 |
| 9 | 4.74230552 | 3.873419686 |

**Supplemental Figure S1**. Ultrametric species tree of *Leptinotarsa* clade with *Anoplophora glabripennis* as an outgroup. Divergence time is stated as millions of years ago (Ma).

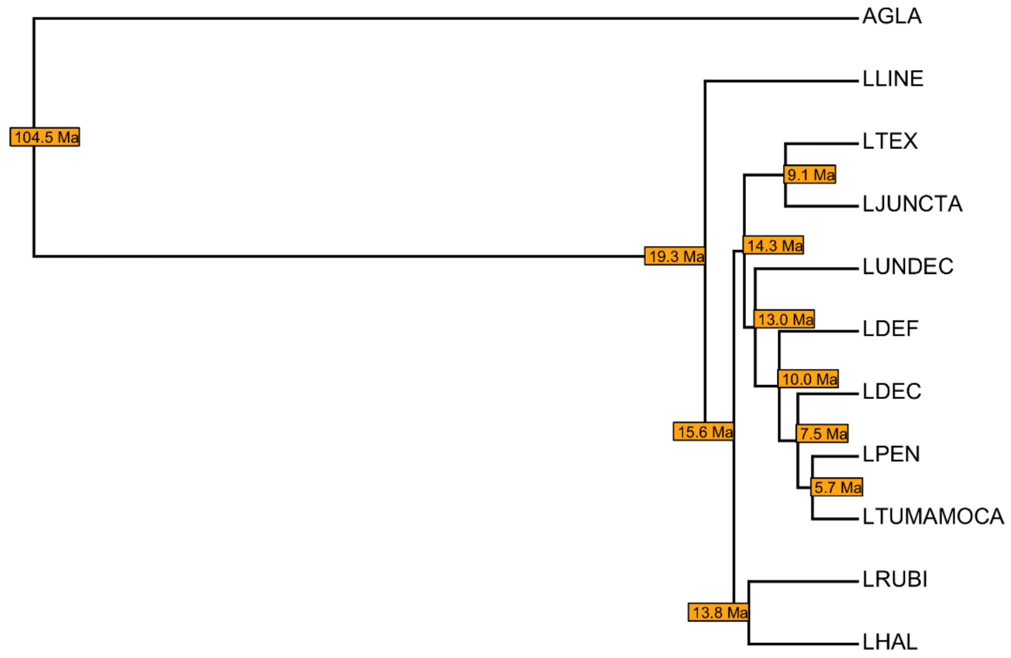


**Supplemental Figure S2.** Boxplot of global heterozygosity in *Leptinotarsa*.

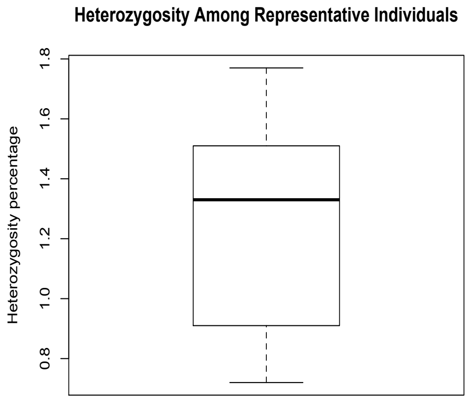

**Supplemental Figure S3.** Venn diagram of GO terms associated with ortholog expansions and contractions in *Leptinotarsa*.

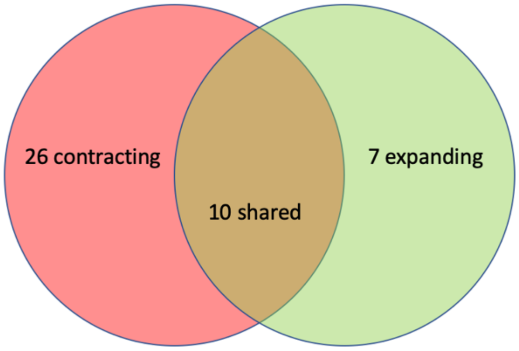


**Supplemental Figure S4**. Number of gene families with significant expansions-contractions within each *Leptinotarsa* species.

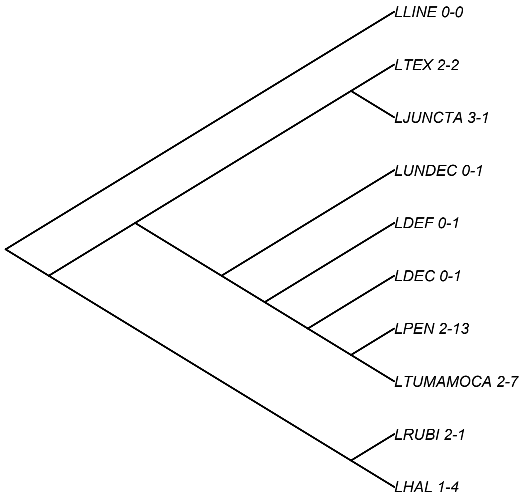


**Supplemental Figure S5**. Number of transposable elements with significant expansions-contractions within each *Leptinotarsa* species.

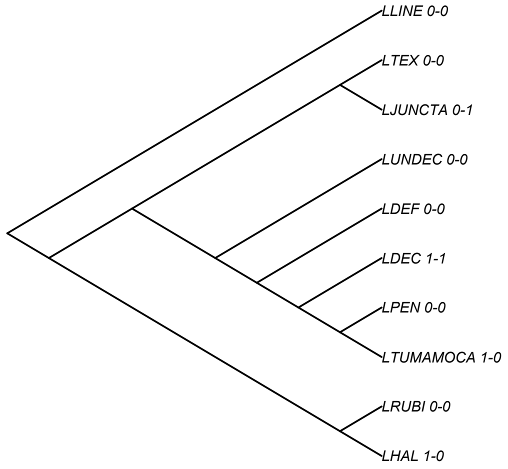

**Supplemental Figure S6.** Heatmap showing the number of shared significant genes among the ten *Leptinotarsa* species compared with the naïve susceptible *L. decemlineata* (CPB) individual. They are arranged from left to right based on genetic distance. With the more recently diverged species from CPB, *L. peninsularis* and *L. tumamoca*, having have fewer significant loci (mean=7.83e+02, stdv= 5.70e+01), than the more distantly related lineages species, *L. undecemlineata*, *L. haldemani*, and *L. lineolata* (mean=1.405e+03, std=9.89e+01). The heatmap shows warmer colors (more shared genes) among the more distantly related taxa, than the more closely related taxa, this is due in part to the greater number of significant genes in these older taxa.  An artifact of this is the increased number of shared loci as one compares longer branches to one another.

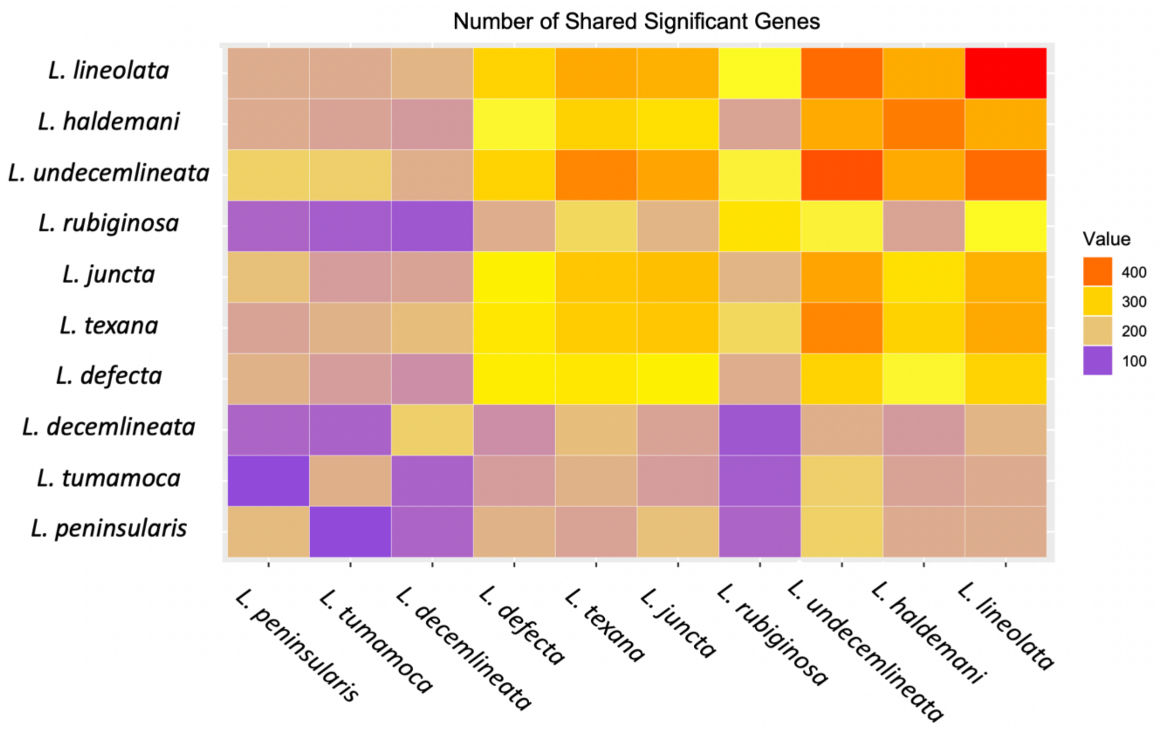

**Supplemental Figure S7**. Venn Diagram representing the unique and shared significant GO terms as well as the unique/shared genes within these GO terms among the three *L. decemlineata* (CPB) genomes.

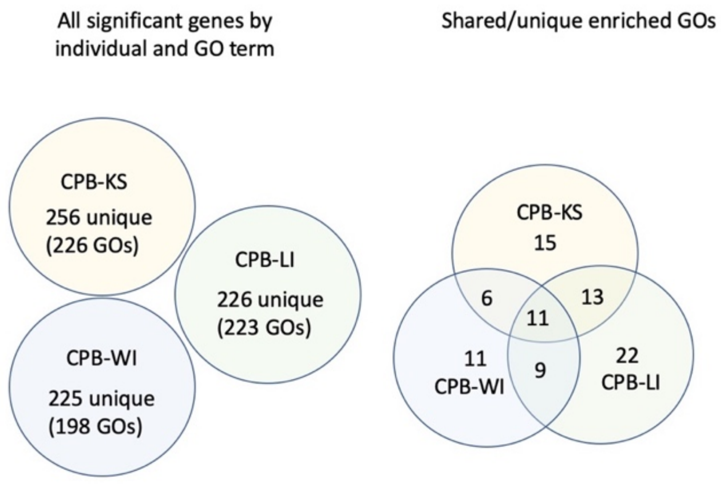


**Supplemental Figure S8.** Boxplots of Standardized PIC values for total and candidate resistant loci under positive selection

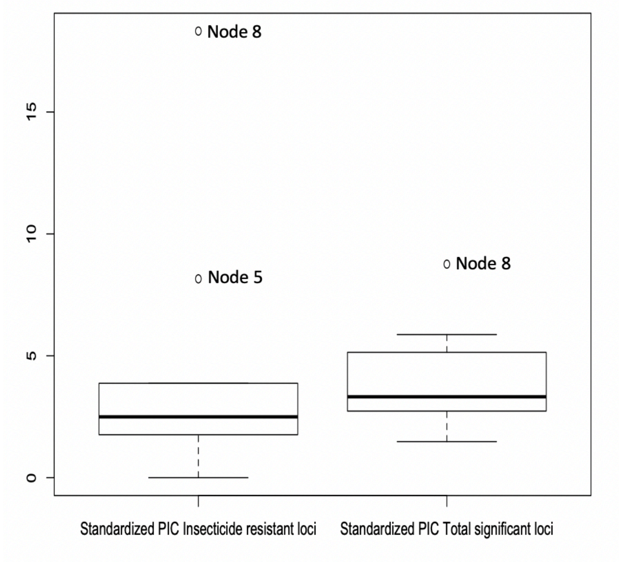
